# Human ventromedial prefrontal cortex lesions enhance expectation-related pain modulation

**DOI:** 10.1101/2021.11.30.470579

**Authors:** JC Motzkin, J Hiser, I Carroll, R Wolf, MK Baskaya, M Koenigs, LY Atlas

**Affiliations:** Department of Neurology, University of California – San Francisco, CA; Department of Anesthesia and Pain Management, University of California – San Francisco, CA; Department of Psychiatry, University of Wisconsin - Madison; Department of Psychology, University of Wisconsin - Madison; Department of Neurological Surgery, University of Wisconsin - Madison; National Center for Complementary and Integrative Health, National Institutes of Health, Bethesda, MD; National Institute of Mental Health, National Institutes of Health, Bethesda, MD; National Institute on Drug Abuse, National Institutes of Health, Baltimore, MD

**Author notes:** Address correspondence to: Lauren Atlas, PhD National Institutes of Health (NCCIH, NIMH, NIDA) 10 Center Drive, Rm 4-1741 Bethesda, MD 20892. These authors contributed equally to this work (M Koenigs, LY Atlas).

**Keywords:** ventromedial, pain, expectations, aversive learning, unpleasantness

## Abstract

Pain is strongly modulated by expectations and beliefs. Research across species indicates that subregions of the ventromedial prefrontal cortex (VMPFC) play a fundamental role in learning and generating predictions about valued outcomes. Consistent with this overarching framework, neuroimaging studies of experimental pain indicate that VMPFC activation tracks expectations of pain relief and statistically mediates expectation-related reductions in responses to painful stimuli across a distributed pain processing network. However, lesion studies in preclinical models and in humans with refractory chronic pain suggest that VMPFC may play a more general role in representing the affective and motivational qualities of pain that contribute to its strong aversive drive. To test whether VMPFC is necessary for pain processing in general, or instead plays a more specific role in the modulation of pain by expectations, we studied responses to experimental heat pain in five adults with bilateral surgical lesions of VMPFC and twenty healthy adults without brain damage.

All participants underwent quantitative sensory testing (QST) to characterize pain sensitivity, followed by a pain expectancy task. Participants were instructed that auditory cues would be followed by heat calibrated to elicit low or high pain. Following a conditioning phase, each cue was intermittently paired with a single temperature calibrated to elicit moderate pain. We compared ratings of moderate heat stimuli and subjective expectancy ratings as a function of cue to evaluate whether VMPFC lesions impact cue-based expectancy and expectancy effects on pain intensity and unpleasantness. We also analyzed QST measures to evaluate whether VMPFC lesions were associated with overall shifts in pain sensitivity.

Compared to adults without brain damage, individuals with VMPFC lesions reported larger differences in expectations as a function of pain-predictive cues, and stronger cue-based modulation of pain ratings, particularly for ratings of pain unpleasantness. There were no group differences in pain sensitivity, nor in the relationship between pain and autonomic arousal, indicating that the impact of VMPFC lesions is specific to expectancy-based modulation of pain unpleasantness.

Our findings suggest that the VMPFC is not essential for basic subjective and physiological responses to painful stimuli. Rather, our findings of significantly enhanced cue-related expectancy effects may suggest that VMPFC plays an important role in updating expectations or integrating sensory information with expectations to guide subjective judgements about pain. Taken together, these results expand our understanding VMPFC’s contribution to pain and highlight the role of VMPFC in integrating cognitive representations with sensory information to yield affective responses.

## Introduction

The role of the frontal lobes in pain has been a focus of considerable attention since early clinical observations of relief from refractory pain following prefrontal lobotomy^1^. Subsequent case series identified midline prefrontal structures and white matter tracts as critical neural substrates of the affective and motivational qualities of pain, with targeted structural lesions resulting in a profound reduction of the “strong aversive drive and negative affect characteristic of pain”^1–3^. These reports are largely consistent with subsequent human neuroimaging studies, which suggest that prefrontal subregions specifically encode the affective qualities of pain, whereas pain location and intensity are represented in more posterior and lateral somatosensory regions^4–7^. However, more recent studies of humans with mixed frontal lesions challenge the specificity of frontal lobe function in pain to affective processes, noting widespread effects on pain detection thresholds, pain tolerance, and ratings of both pain intensity and pain unpleasantness^8–10^.

More recently, functional neuroimaging studies have implicated subregions of human ventromedial prefrontal cortex (VMPFC) in the modulation of pain by expectations. Acute pain is highly susceptible to modification by expectations and by predictive cues signaling pain or relief^11–14^. In prior work, the magnitude of expectation-related activation in VMPFC is reliably associated with the strength of expectancy effects on subjective pain ratings^15, 16^. Further, activity in this region has been shown to statistically mediate the effects of predictive cues on pain-related activity across a distributed pain processing network, which in turn give rise to subjective pain^17^. Preclinical models showing a loss of cue-based pain modulation following inactivation of homologous regions of infralimbic cortex offer further support for a specific modulatory function of VMPFC in the context of pain^18^.

Theoretical accounts of VMPFC function across species and experimental contexts may provide a more powerful way to understand the varied roles ascribed to this region in the context of pain (for review, see ^19, 20^). One overarching view of VMPFC function is that it plays a central role in the formation of schemas^21, 22^ and cognitive maps^23–25^ in response to latent or inferred rules^26, 27^ from which to guide learning, decision-making, and valuation. As the experience of pain is highly sensitive to predictions and expectations^28–30^, VMPFC might play a central role in shaping pain, and more specifically, pain affect, through the generation of valuation and affective meaning^31^. Importantly, this model is generally consistent with prior human lesion work showing that VMPFC damage attenuates physiological arousal during aversive learning^32^, and that altered anticipatory responses may themselves impair decision making and social behavior^33–36^.

Together, these accounts offer support for an integratory function of VMPFC, in which cognitive biases, somatosensory cues, and autonomic signals are combined to update judgements of subjective value in the context of pain^31, 37^.

Despite the substantial body of clinical and experimental evidence implicating VMPFC as a critical link between cognitive state and the affective-motivational features of pain, there are as-yet no systematic studies in humans investigating the effects of focal damage to VMPFC on pain perception and modulation. Here, we use a well-validated experimental procedure (Figure 1) ^17, 38–40^ in a population of five neurosurgical patients with focal, stable, bilateral VMPFC lesions and twenty healthy adult comparison subjects (HC) to test three central hypotheses of VMPFC function in pain; namely that VMPFC lesions would (1) increase subjective pain thresholds and pain tolerance, (2) reduce the impact of pain-predictive cues on expectations about pain, subjective pain ratings, and autonomic responses to painful stimuli, and (3) reduce the typical associations between subjective pain ratings and autonomic responses to painful stimuli.

**Figure 1.**
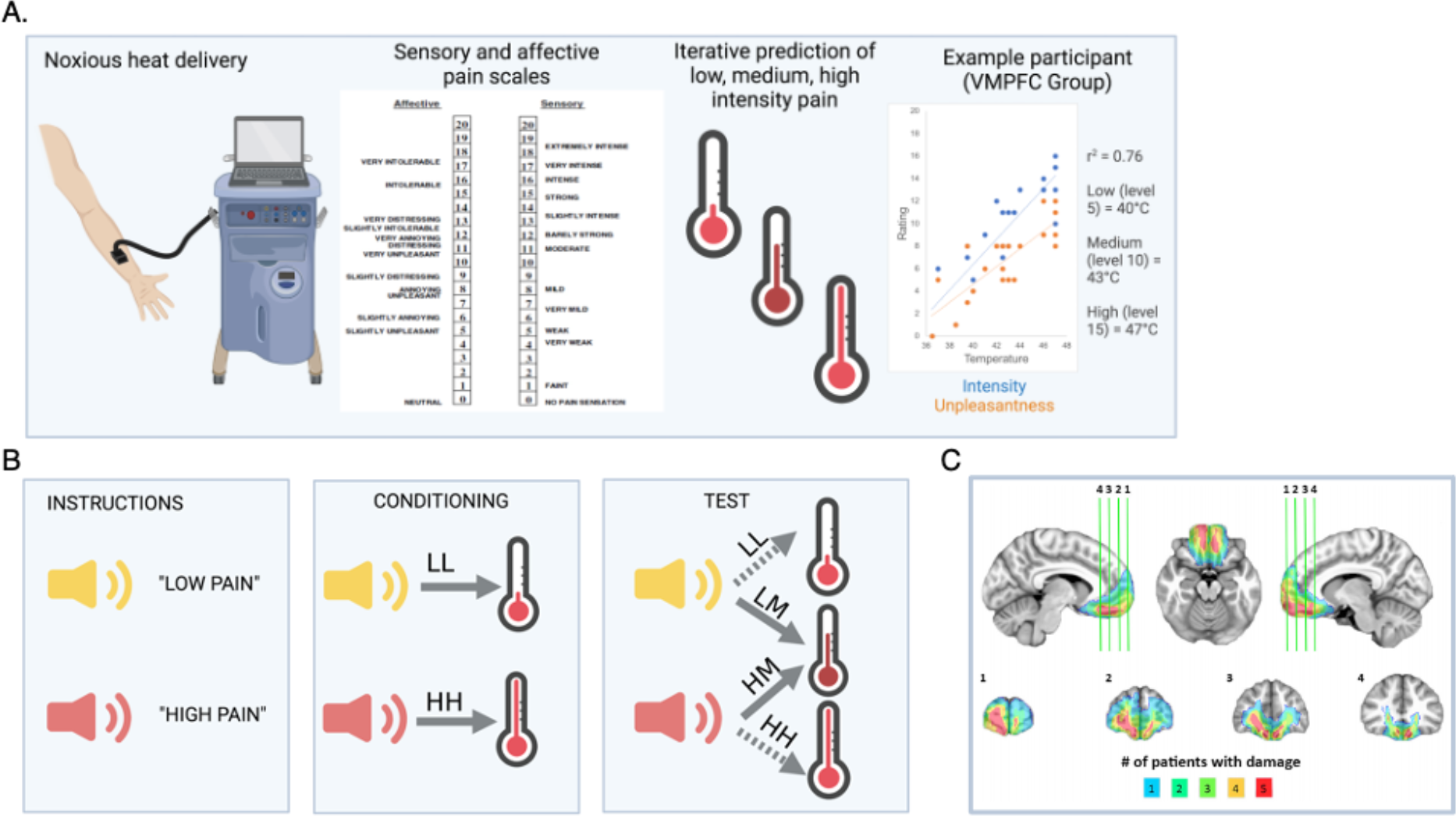
Experimental design and VMPFC lesion overlap. **(A)** Graphical representation of the adaptive pain calibration procedure including the pain stimulation device and thermode (left), representative scales of pain unpleasantness (affective) and intensity (sensory) with verbal groundings of numerical ratings (center), and the calibration procedure used to identify individual temperatures expected to elicit low, medium, or high levels of pain (right). Example calibration graph shows ratings of intensity (blue) and unpleasantness (orange). For this patient, calibrated low, moderate, and high pain levels are 40°C, 43°C, and 47°C, respectively. **(B)** Graphical representation of the pain expectancy task showing initial presentation of auditory cues (left), followed by two blocks of conditioning trials in which cue is reliably followed by predicted level of heat (center), and six blocks of test trials (right) in which low- or high-pain cues are followed either by predicted low and high temperatures (LL and HH trials) or a temperature calibrated to elicit moderate pain (LM and HM trials). LL=Low Pain Cue+Low Heat, LM=Low Pain Cue+ Moderate Heat, HM=High Pain Cue +Moderate Heat. HH=High Pain Cue + High Heat. (**C)** Lesion overlap map depicting lesion extent across the VMPFC lesion group. All subjects have bilateral lesions involving the ventral third of the medial frontal cortex and medial third of the orbitofrontal cortex. Colors indicate number of patients with damage in a particular region.

## Materials and methods

### Participants

All experimental procedures were approved by the University of Wisconsin – Madison Institutional Review Board (IRB). The target lesion group consisted of five adult neurosurgical patients with focal, bilateral parenchymal changes largely confined to the VMPFC, defined as the medial one-third of the orbital surface and the ventral one-third of the medial surface of prefrontal cortex (see Figure 1). All 5 patients acquired brain lesions as a result of meningioma growth and ultimate resection via anterior skull base craniotomy. All lesions reflect a combination of direct parenchymal damage from long standing meningiomas, vasogenic edema, and definitive surgical resection. Initial clinical presentations included subtle or obvious personality changes over at least several months preceding surgery. On postsurgical MRI, though evidence of vasogenic edema had largely resolved, there were persistent T2-weighted signal changes, consistent with gliosis, in VMPFC. All patients were adults at the time of surgical resection and all experimental data were collected at least three months after surgery. At the time of testing, all patients had focal, stable resection cavities on MRI and were free of dementia and substance use disorder. All patients had no history of pain disorder, myocardial infarction, presence of implanted cardiac device, or current use of analgesic medication.

Twenty healthy adults with no history of brain injury, myocardial infarction, presence of implanted cardiac device, neurological or psychiatric illness, pain disorder, or current use of psychoactive or analgesic medication were recruited as a healthy comparison (HC) group. Demographic and neuropsychological data for both groups are summarized in Table 1.

**Table 1.**
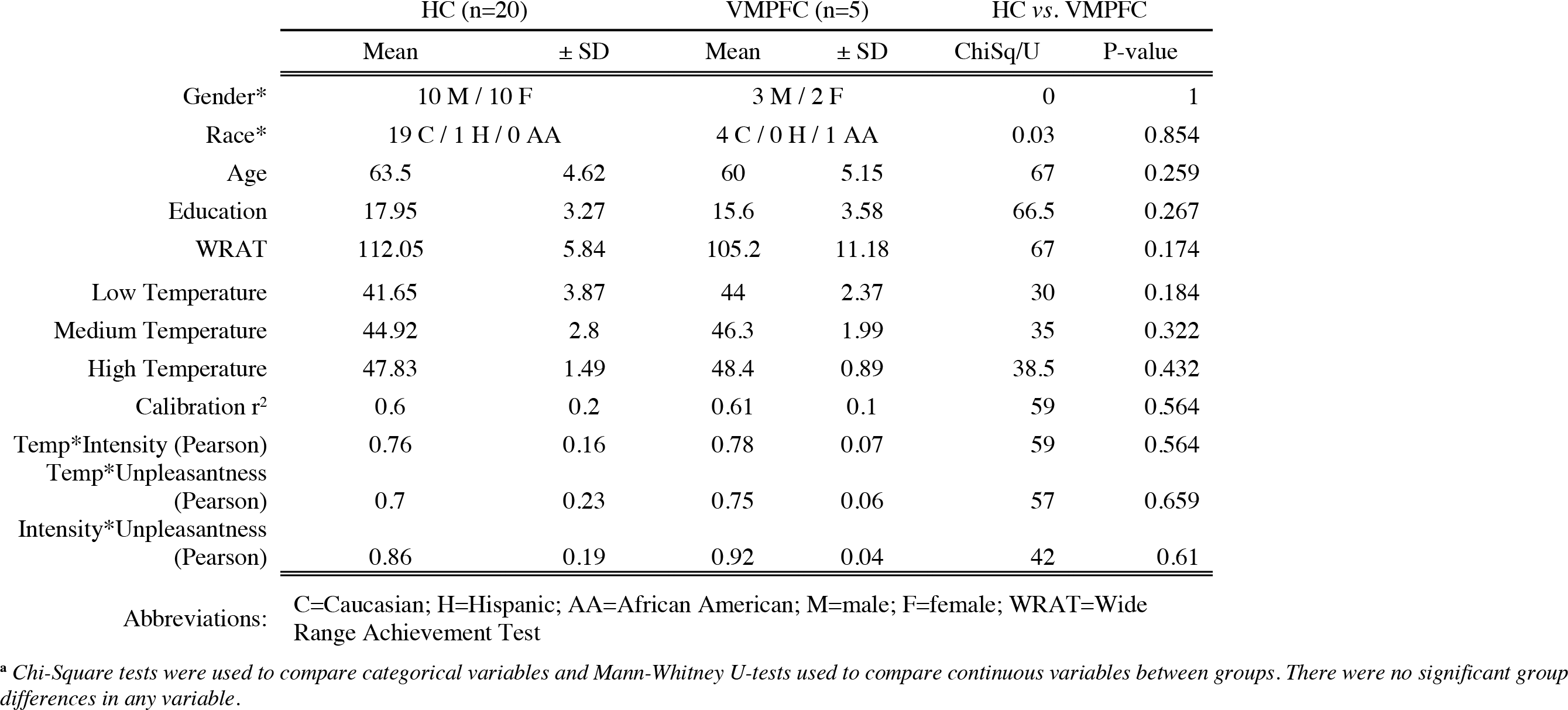
Demographics and Pain Calibration Data for HC and VMPFC Groups^a^

### Lesion segmentation and image normalization

Structural MRIs for VMPFC patients were obtained at least three months after surgery. Individual VMPFC lesions were visually identified and manually segmented on T1-weighted images. Lesion boundaries were drawn to include areas with gross tissue damage or abnormal signal characteristics on T1 or T2 FLAIR images. T1-weighted images were skullstripped and diffeomorphically aligned to the Montreal Neurological Institute (MNI) coordinate system using a symmetric normalization algorithm with constrained cost-function masking to prevent warping of tissue within the lesion mask^41, 42^. The lesion overlap map (Figure 1C) was created by computing the sum of aligned binary lesion masks for all five VMPFC patients.

## Materials and Procedure

### Overview

To test the three study hypotheses, we performed an adaptive thermal pain calibration procedure^17, 43^ to identify individually-calibrated temperatures expected to elicit low, medium, or high levels of pain, followed by a pain expectancy task^17^ in which Low or High pain-predictive cues were conditioned with low or high temperatures, respectively, and then paired with medium temperatures to isolate expectancy effects on pain and autonomic responses. Participants provided ratings of pain intensity and pain unpleasantness for each stimulus and autonomic indices of physiological arousal (including skin conductance response [SCR] and heart rate [HR]) were recorded throughout.

### Stimuli and apparatus

Thermal stimuli were delivered to the volar surface of the left forearm using a 30x30 mm Peltier thermode (Medoc, Inc). Each stimulus lasted 11 seconds, with 2-second ramp-up and ramp-down from a 32°C baseline and 7 seconds at target temperature. Target temperatures were individually calibrated for each participant (see “Pain calibration and threshold procedure”). We did not apply any temperatures above 49°C to avoid skin damage.

Following heat offset, participants separately rated pain intensity and pain unpleasantness for each stimulus using scales adapted from Petzke *et al* ^44^, with standardized verbal descriptors displayed alongside numerical ratings (See Figure 1A). The intensity scale was a 21-box Likert scale ranging from 0 (no pain sensation) to 20 (unbearable pain). Prior to calibration, participants were informed that an intensity rating of 5 on this scale should indicate the temperature at which heat is first perceived as painful (i.e., pain threshold) and that a rating of 15 should indicate the maximum temperature they would be willing to tolerate during the experiment (i.e., pain tolerance). The unpleasantness scale was a 21-box Likert scale range from 0 (neutral) to 20 (very intolerable). Participants were instructed that intensity ratings should reflect how the pain “feels to you” whereas unpleasantness ratings should reflect how the pain “makes you feel”. Verbal descriptors were present on rating scales throughout the experiment. During the study, auditory cue presentations, thermal stimulus triggers, and collection of pain ratings were implemented using E-prime software (Psychology Software Tools, Pittsburgh, PA).

Heart rate data were acquired at 2000 Hz with a BioPac photoplethysmograph (Biopac Inc), affixed to the left index finger throughout the experiment. Skin conductance responses (SCRs) were collected at 10 Hz according to standard guidelines^45, 46^ using 2 shielded Ag-AgCl electrodes prepared with isotonic paste and affixed to the second and third digits of the participants’ left hand. All physiologic data was collected using the BioPac MP150 system.

### Procedure

#### Pain calibration and threshold procedure

Before introducing the expectancy manipulation, all subjects first underwent a quantitative sensory testing (QST) calibration procedure using four sites on the volar surface of the left forearm (see Figure 1A). We used an adaptive staircase procedure, described in detail in previous work^17, 43, 47, 48^, to estimate the dose-response curve of the relationship between applied thermal stimulation and reported pain intensity (slope, intercept, r^2^) for each participant. In the present study, participants rated both subjective intensity and unpleasantness after heat offset using the corresponding 21-item Likert scales. We used linear regression to iteratively fit pain intensity ratings as a function of temperature for each participant in order to derive temperatures predicted to elicit Low pain (i.e. pain threshold; intensity rating of 5), Moderate pain (intensity rating of 10), and High pain (i.e. maximum tolerance; intensity rating of 15), which were then used in the main expectancy experiment. Only participants who had an r^2^ > 0.4 at the end of the calibration procedure were included in the experimental task analyses, consistent with prior work.^17^ Three healthy volunteer participants were excluded from the expectancy task on the basis of low r^2^ values, such that sample size in the HC group for calibration analyses was 20 and for expectancy task analyses was 17. All participants in the VMPFC group had r^2^ > 0.4 (see Table 1) and were thus included in all analyses.

#### Expectancy instruction and training task

All participants were explicitly informed of cue-pain contingencies prior to any experimental pairings between cues and heat (see Figure 1B). Cues were 2-second auditory tones (a cymbal or a cowbell). Cue-outcome contingencies were counterbalanced across participants. Participants were instructed that one tone would predict low pain and the other tone would predict high pain. Each tone was played along with the instruction “This is your [low/high] pain tone” three times before beginning the training task. Following instructions, participants completed a discrimination task consisting of 10 trials in which they had to correctly identify the expected level of pain following each cue. Predictive cues were presented in random order, and participants used the L key to identify Low Pain cues, and the H key to identify High Pain cues. Participants were required to successfully identify at least 90% of trials to proceed to the experimental portion of the study, consistent with previous work^17^. No participants were excluded on this basis.

#### Task design and trial structure

After calibration and training, experimental data were acquired across 8 blocks (8 trials/block, 64 total trials per participant), using a task adapted from Atlas *et al*.^17^ The thermode was placed on a different skin site for each block, with two total blocks per skin site. On each trial, a two-second pain-predictive cue (Low Pain Cue or High Pain Cue) was followed by a four-second anticipatory interval during which a fixation cross was presented on the screen. Thermal stimulation was then delivered via the thermode for eleven seconds (2s ramp-up from 32°C baseline, 7s at peak destination temperature, 2s return to baseline), with destination temperatures determined based on each participant’s calibration. Six seconds after heat offset, participants were prompted to record pain ratings of pain intensity and pain unpleasantness.

There were four types of trials in the expectancy task (see Figure 1B). On “Low Pain Cue + Low Heat” (LL) trials, Low Pain cues were followed by stimulation calibrated to elicit an intensity rating of 5 based on the pain calibration procedure. On “High Pain Cue + High Heat” (HH) trials, High Pain cues were followed by stimulation calibrated to elicit intensity ratings of 15. On “Low Pain Cue + Moderate Heat” (LM) trials and “High Pain Cue + Moderate Heat” (HM) trials, Low or High Pain cues, respectively, were followed by stimulation calibrated to elicit intensity ratings of 10 (Moderate Pain). Thus, consistent with Atlas *et al.,*^17^ applied temperatures were identical in the critical LM and HM trials but the preceding cues (falsely) indicated high or low intensity stimulation based on prior explicit instruction and reinforcement through conditioning.

Participants first experienced two blocks (16 trials) of trials evenly divided between LL and HH trials, in pseudorandom order (see Figure 1B). This conditioning phase served primarily to reinforce the instructed cue-outcome relationships through associative learning. These were followed by six blocks of trials equally divided between LL, HH, LM, and HM trials, presented in pseudorandom order and counterbalanced across participants. Participants were not informed that moderate heat stimulation would be delivered, and thus the comparison of HM and LM trials provides a measure of how cue-based expectations shape pain. For additional details on task design, please see Atlas *et al*.^17^

#### Expectancy ratings

Before the first run and after each run, participants heard the Low Pain and High Pain cues and were asked, after each tone, “When you hear this tone, how much pain do you expect?” Participants then rated both expected intensity and unpleasantness. The first such ratings were made prior to any pairing between predictive cue and thermal stimulation, thus providing a measure of cue effects on pain expectancy from instructions alone, unaffected by associative learning. The remaining eight ratings served as a manipulation check, allowing us to measure conscious cue-related expectancies and to test whether expectancies changed over the course of the experiment.

### Data processing and analysis

#### Heart rate analysis

To assess cardiac responses to thermal stimuli, we computed trial-wise estimates of heart rate change for each subject^49^. Peaks in the photoplethysmograph tracing, corresponding to cardiac R-waves in the QRS complex, were identified using in-house interactive beat detection software. Trials with ectopic beats, missed beats, or periods of noisy signal (where beat detection failed), were excluded from further analysis. R-R intervals were transformed into heart rate in beats per minute, in 500ms bins. Changes in heart rate were determined by subtracting the mean heart rate for 2s preceding each thermal stimulation from the mean heart rate between 1 and 14 seconds after thermal stimulus onset. Mean heart rate change was computed separately for each trial.

#### Skin conductance analysis

Skin conductance data were analyzed in Matlab (Mathworks, Natick, MA) using Ledalab (http://www.ledalab.de). Specifically, using Continuous Decomposition Analysis^50^, the skin conductance data were deconvolved into phasic and tonic drivers of the skin conductance response (SCR). The response window for skin conductance fluctuations to be regarded as stimulus related was defined as 1-14 seconds following the onset of the thermal stimulus. To correct for the skewed distribution of the skin conductance data, we square-root transformed SCR values ^45, 51^. We analyzed both square-root transformed SCR and z-scored SCR within participants, which account for variations in amplitude between groups. We focus on the sum of amplitudes from Continuous Decomposition Analysis as a measure of phasic heat-induced SCR in the main manuscript and report additional phasic and tonic outcome measures in Supplementary Materials.

#### Statistical Analysis

To test our first hypothesis, that VMPFC lesions would increase pain thresholds and pain tolerance, we evaluated pain sensitivity measures derived from the adaptive staircase calibration procedure. We used ANOVAs with Type III Sums of Squares, implemented using the car package^52^ in R statistical software suite (Version 4.0.3)^53^ to evaluate whether groups differed in the individually calibrated low, medium, and high temperatures that were applied during the subsequent experiment. We also used linear mixed effects models to test for group differences in the effects of temperature on pain intensity and pain unpleasantness ratings across all trials during calibration. Details regarding linear mixed model specification are below. As a final test of Hypothesis 1, we performed an identical linear mixed effects analysis using data from the subsequent experiment to evaluate, across all 64 trials, whether VMPFC lesions were associated with altered temperature-rating relationships. Group differences in demographic variables and post-hoc comparisons of pain thresholds were assessed using Chi-square and Mann-Whitney-Wilcoxon tests implemented in R.

To test our second hypothesis, that VMPFC lesions would reduce the impact of pain-predictive cues on expectations, subjective pain ratings, and autonomic responses to painful stimuli, we examined effects of Group and Predictive Cue (Low or High) on 1) expectancy ratings collected before each block of the experiment; 2) pain ratings on critical medium heat trials; and 3) trial-wise estimates of autonomic reactivity to the medium heat trials. We used t-tests and ANOVAs to examine group differences in subjective expectancy ratings following instructions and after each block in the test phase. Expectancy ratings were analyzed as difference scores (i.e., High Pain Cue expectation – Low Pain Cue expectation) and as a function of Cue (see Supplement). To test the effects of expectancies on pain and autonomic responses to heat, we evaluated the interaction between Group and Cue on responses to medium heat trials using linear mixed models, as described below.

Finally, we tested our third hypothesis, that VMPFC lesions would reduce the typical associations between subjective pain ratings and autonomic responses to painful stimuli. We performed linear mixed effects analyses with separate trial-wise regressors for SCR or HR, mean centered for each subject. To investigate group differences in the trial-by-trial correlation between ratings and autonomic variables, we examined the interaction between group and mean-centered ratings on each autonomic variable in the subset of medium temperature trials.

All linear mixed models were implemented using the lme4 package^54^ in the R statistical software suite (Version 4.0.2), and confirmed with the package nlme^55^ which allowed us to account for autoregression, and with Bayesian models implemented in brms^56^ that allowed us to provide posterior estimates on the magnitude of the effects. We used the package bayestestR^57^ to evaluate posteriors and evaluate practical significance, i.e. our ability to accept or reject the null hypothesis, as discussed in Makowski *et al.*^57, 58^ For each analysis, we mean centered all predictors (Temperature, Cue, and Trial) and modeled Group as a mean-centered variable to aid with interpretation of the intercept. Effects of time (Trial, Block) were treated as fixed, while effects of Temperature and Cue were modeled with random slopes in all models unless there were issues with convergence. See Supplementary Methods for details of model specification, reporting, and evaluation of significance.

### Data availability

Raw data were generated at University of Wisconsin - Madison. Derived data supporting the findings of this study are available from the corresponding author on request.

## Results

### Individuals with VMPFC lesions have similar pain sensitivity to healthy comparison participants based on quantitative sensory testing

We first analyzed data from the adaptive QST calibration to test the hypothesis that VMPFC lesions would impact pain sensitivity (i.e., pain thresholds, pain tolerance, and the relationship between stimulus temperature and ratings). Group means and comparisons are reported in Table 1. There were no differences between groups in the association between temperature and intensity ratings during calibration as measured by r^2^ or Pearson’s r, nor in the association between temperature and unpleasantness ratings, nor between intensity and unpleasantness ratings (see Table 1), indicating that subjective judgments about pain track stimulus temperature similarly between groups.

To further assess group differences in the relationship between temperature and subjective pain, we evaluated ratings of pain intensity and unpleasantness across all trials during the calibration as a function of group and temperature using linear mixed models (see Figure 2 and Supplementary Table 1). As expected, we observed a significant main effect of temperature for both types of ratings (intensity: !LMER=1.17, p < .001; unpleasantness: !LMER=0.84, p < .001) that were practically significant (0% of posterior estimates in ROPE), indicating that higher temperatures were rated as more intense and unpleasant. We did not observe any main effect of Group nor any Group x temperature interactions for either measure (all p’s > 0.1; see Supplementary Table S1), and >99% of posterior estimates for the Group x Temperature interactions fell within the ROPE, supporting the null hypothesis of no group differences in temperature effects on pain intensity or unpleasantness. Thus, the VMPFC lesion group showed similar associations between variations in temperature and subjective pain during the adaptive staircase calibration.

**Figure 2.**
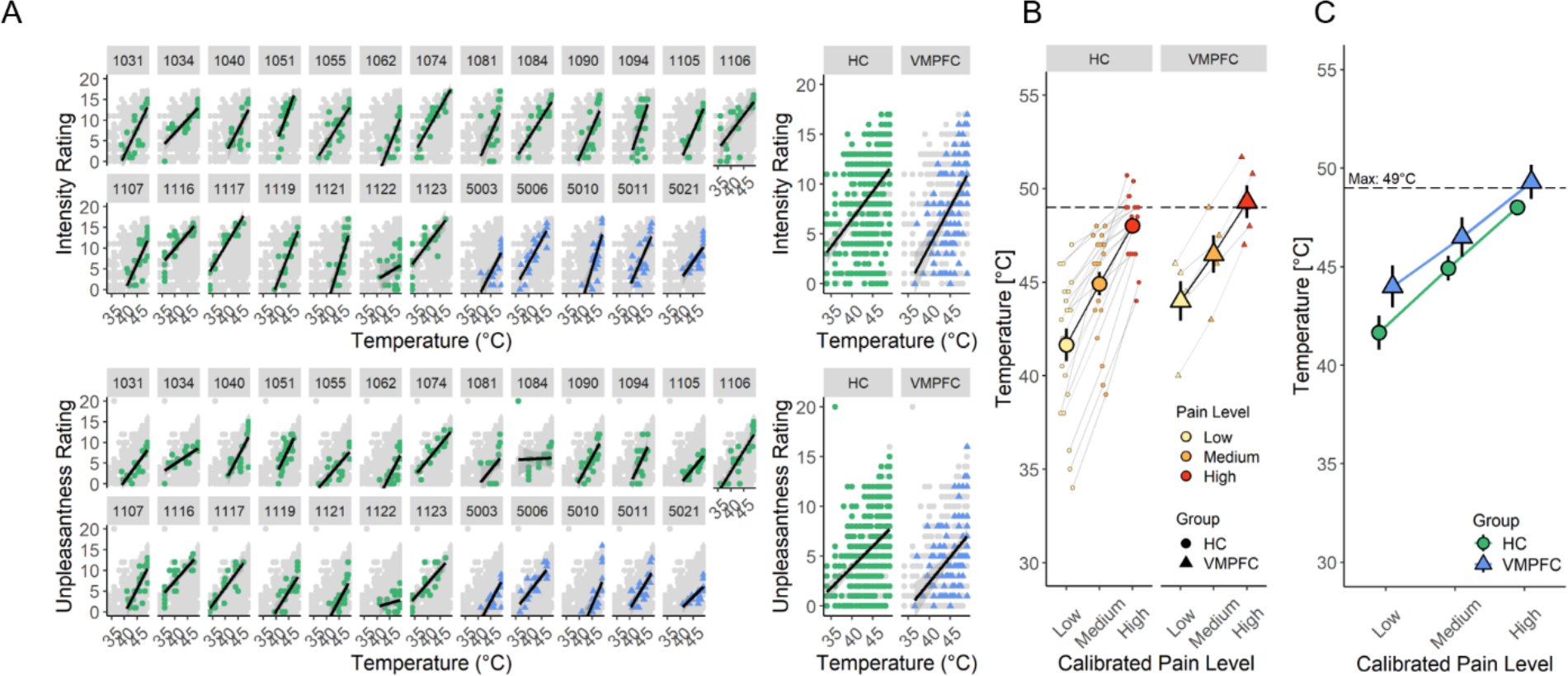
Adaptive staircase calibration and temperatures used during the task. **(A)** Scatterplots depicting correlation between applied temperatures and ratings of intensity (top row) and unpleasantness (bottom row) for individual subjects (left) and across the entire group (right). HC subjects (green circles) and VMPFC subjects (blue triangles) have similar temperature-rating curves. **(B)** Line graphs depicting correlation between estimated pain level (low, medium, high) and temperature for HC (left) and VMPFC (right) groups. Group means are shown in bold over individual subject values. **(C)** Summary of group means for HC (green circles) and VMPFC (blue triangles) across all 3 temperatures. Though temperatures appear higher in the VMPFC group, this difference did not meet criteria for statistical significance.

Finally, we tested whether groups differed in the individual temperatures calibrated to elicit low pain (i.e., pain threshold), moderate pain, and high pain (i.e., pain tolerance) for the subsequent experiment (Table 1, Figure 2B). A 2 (Group) by 3 (Pain Level) mixed ANOVA revealed the expected significant main effect of pain level (F(2, 69)=64.06, p<0.001). Although estimates for the VMPFC group were between 1-3℃ higher than the HC group on average for each of the three estimated temperatures, there were no main effects of Group, nor Group x Pain Level interactions (all p’s > 0.1), indicating that pain sensitivity was not significantly different between groups during calibration. Post-hoc non-parametric comparisons at each estimated temperature level were similarly non-significant (Table 2, all p’s > 0.1). Thus, contrary to our first hypothesis, analyses of calibration data indicate that pain sensitivity, pain thresholds, and pain tolerance are not affected in individuals with VMPFC lesions. Importantly, this finding indicates that any observed differences between groups that emerge during the expectancy task cannot be explained by group differences in pain sensitivity or applied temperatures, allowing us to test our hypotheses about the effects of VMPFC lesions on expectations and expectancy-based pain modulation.

**Table 2.**
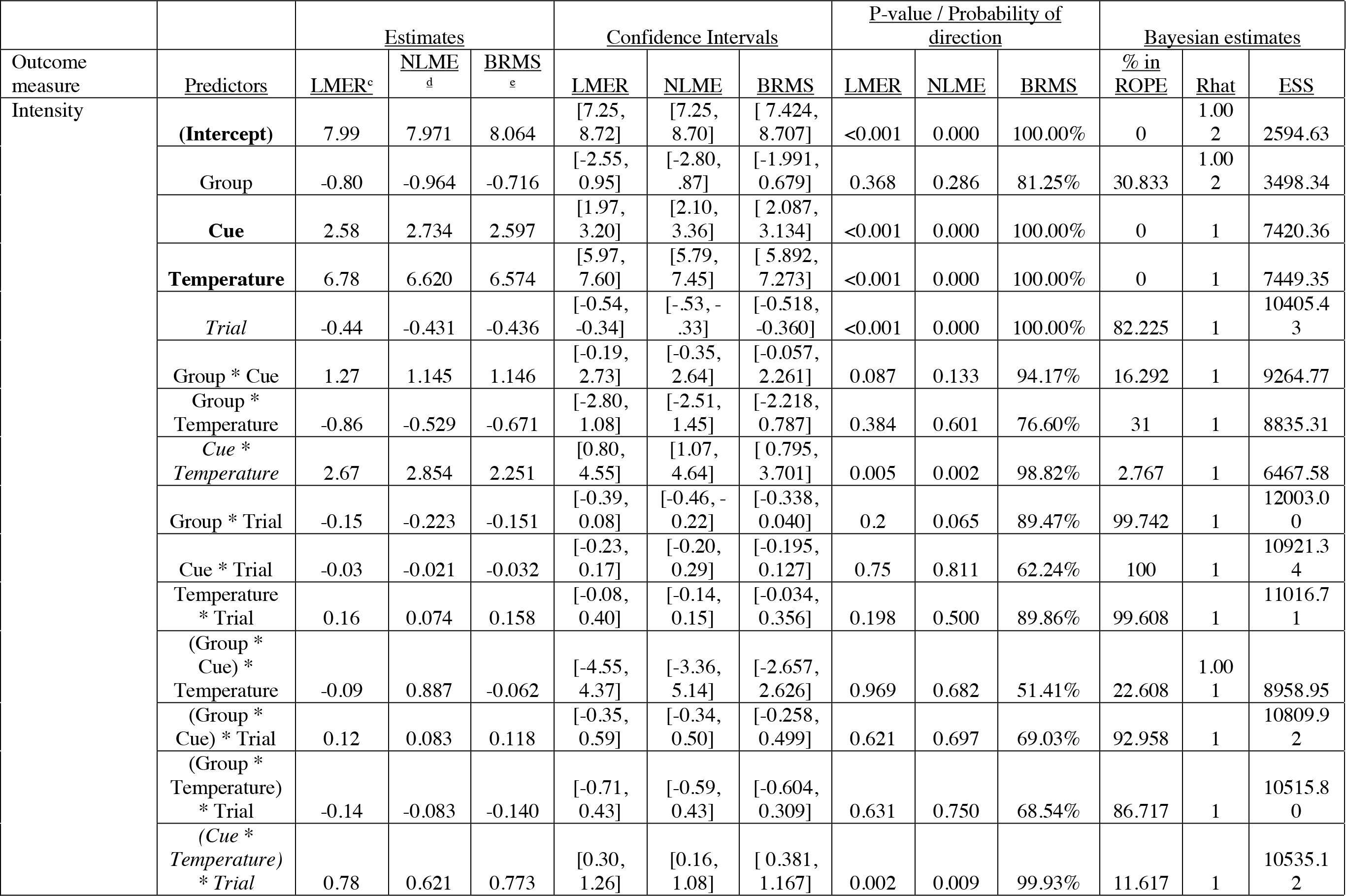
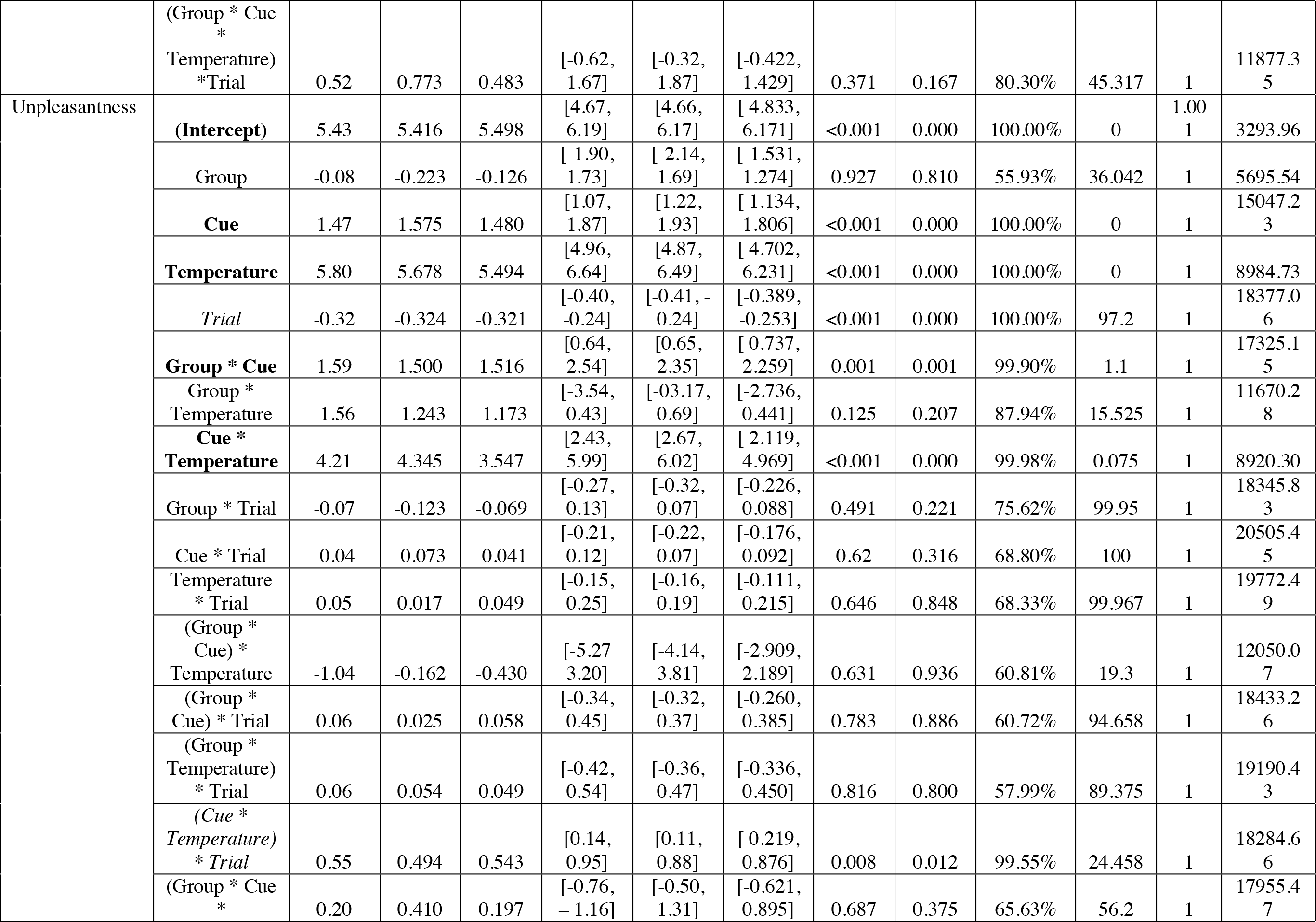
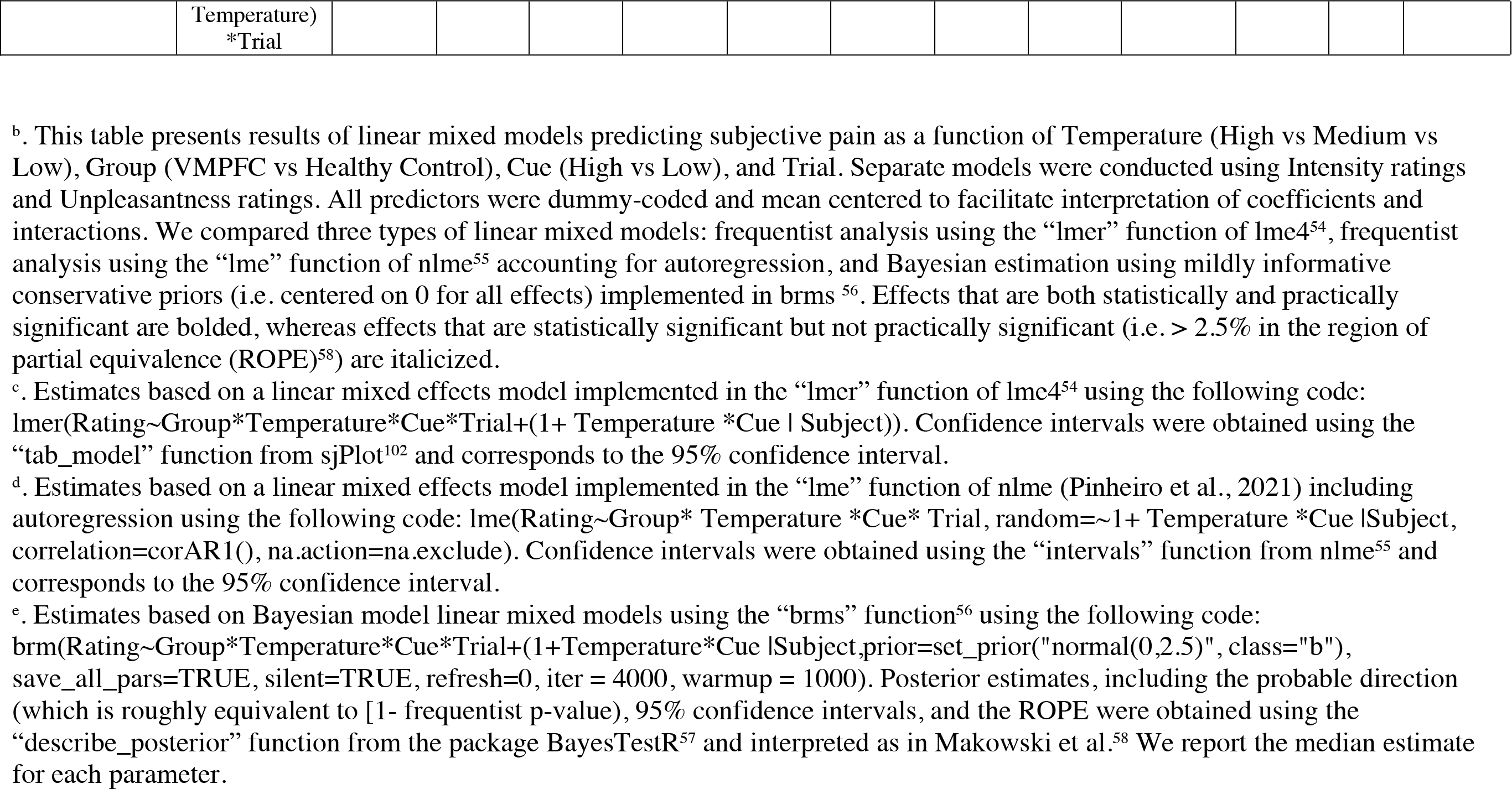
Temperature effects on pain during the main experiment.^b^

### Temperature effects on pain do not differ by group

To provide a secondary test of Hypothesis 1, we evaluated pain as a function of temperature during the experimental phase of the study to determine whether VMPFC lesions alter pain-related responses across all trials regardless of expectancy. Complete results are reported in Table 2, indicating strong main effects of Temperature, Cue, and Trial, regardless of analytic approach (all p’s < 0.001). Effects of Temperature and Cue were practically significant (0% in ROPE). Consistent with the fact that noxious stimuli were individually calibrated prior to the experimental phase, we did not observe a main effect of Group, nor any interactions between Group and Temperature (all p’s > 0.1) for either type of pain rating. The only difference between groups that was practically significant was a Group x Cue interaction in unpleasantness (all p’s < .001; 1.1% in ROPE) suggesting that predictive cues influenced unpleasantness and that these effects may have differed by group. We therefore turned to analyses of self-reported expectations and pain ratings on the critical medium heat trials to test our second hypothesis, that VMPFC lesions alter expectancy-based pain modulation.

### VMPFC lesions are associated with larger differences in expected pain based on pain- predictive cues

To test our hypothesis that individuals with VMPFC lesions would report altered explicit cue-based expectations, we analyzed group differences in the acquisition and evolution of explicit cue-based expectancies across the task. We analyzed expectancy ratings collected before each block of the task. We computed the difference in expected pain between cues (i.e., High Pain Cue expectation – Low Pain Cue expectation) to derive an index of cue-based expectations. The first rating was collected immediately after introducing the instructed contingencies, before any pairing between cues and heat stimuli. Prior to learning, individuals with VMPFC lesions reported larger differences in expected intensity than individuals without lesions (t(14.436) = - 2.57, p = 0.022; see Figure 3); there were no group differences in expected unpleasantness prior to conditioning (p = .27). There were no group differences in expectancy ratings during the conditioning phase (all p’s > 0.1; see Figure 3). During the test phase, patients with VMPFC lesions reported larger cue-based differences in expected unpleasantness (F(1,20) = 8.475, p = .009) and marginally larger differences in expected intensity (F(1,20) = 4.15, p = .055). Findings based on difference scores were confirmed with analyses separated by cue type using both ANOVAs and linear mixed models (see Supplementary Results). Linear mixed models indicated that the Group x Cue interaction on expected unpleasantness was practically significant (all p’s < 0.01; 2.28% in ROPE; see Supplementary Table S2), while the Group x Cue interaction on expected intensity was of undecided significance (all p’s < 0.05; 8.61% in ROPE). Thus, VMPFC lesions influenced expectations for both Low and High pain cues, and groups did not differ significantly in response to either cue alone (see Figure 3).

**Figure 3.**
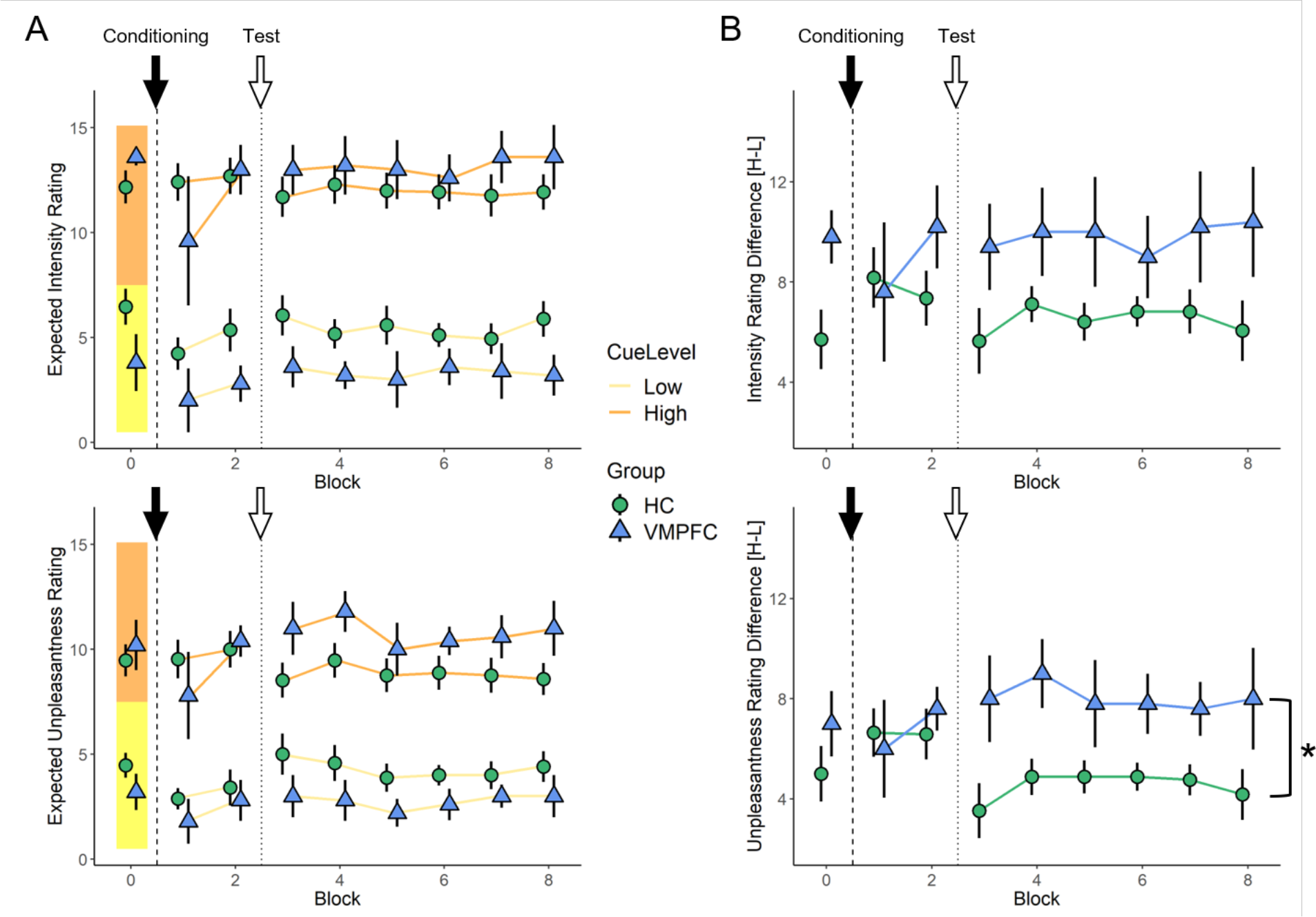
Behavioral Results: Expectancy ratings. **(A)** Expected pain intensity (top) and unpleasantness (bottom) after hearing High (orange) or Low (yellow) pain predictive cues. **(B)** Difference in expected pain intensity (top) and unpleasantness (bottom) between High and Low pain predictive cues for HC (green circles) and VMPFC (blue triangles) groups. Block 0 ratings are acquired at baseline, before any pairings between cue and heat. Black arrows and dashed lines indicate the start of the conditioning phase of the task and white arrows and dotted lines indicate the introduction of moderate temperature trials, signaling the beginning of the test phase of the task. The VMPFC group maintained significantly greater differences in expected pain unpleasantness during the test phase (*p<0.001), driven by similar cue effects for both low and high levels of expected pain.

Thus, in contrast to our second hypothesis that VMPFC lesions would impair the acquisition of cue-related expectations, we found that lesions actually *enhance* instructed expectations about predictive cues. Furthermore, though differences are present even based on instruction alone, experiential learning (i.e., pairings between instructions and reinforcement) serves to enhance group differences, particularly in expected unpleasantness.

### Individuals with VMPFC lesions report larger cue effects on pain unpleasantness

As a second test of hypothesis 2, we asked whether VMPFC lesions reduce the impact of cue-based expectations on subjective pain ratings. We analyzed data from the critical moderate heat trials crossed with Low and High pain-predictive cues (i.e., LM and HM conditions) to isolate expectancy effects at a constant level of thermal stimulation. Across analytic approaches (see Table 3), there was a practically significant main effect of Cue on both types of ratings (all p’s < .001; 0% in ROPE), and ratings generally decreased over time, although the effect of time was of undecided significance (see Table 3). Contrary to our hypothesis, but in line with our findings based on explicit expectancy ratings, individuals with VMPFC lesions showed *stronger* effects of pain-predictive cues on ratings of unpleasantness, as indexed by a practically significant Group x Cue interaction (all p’s < 0.001; 0.5% in ROPE). We also observed Group x Cue interactions in pain intensity in the same direction, although this term was marginal in frequentist analyses and of undecided significance based on Bayesian models (see Table 3).

**Table 3.** Pain-predictive cue effects on pain and SCR evoked by medium heat.^f^

|  |  | <u>Estimates</u> |  |  | <u>Confidence Intervals</u> |  |  | <u>P-values / Probability of direction</u> |  |  | <u>Bayesian estimates</u> |  |  |
| --- | --- | --- | --- | --- | --- | --- | --- | --- | --- | --- | --- | --- | --- |
| <u>Outcome measure</u> | <u>Predictors</u> | <u>LME R</u> | <u>NLM E</u> | <u>BRM S</u> | <u>LMER</u> | <u>NLM E</u> | <u>BRMS</u> | <u>LMER</u> | <u>NLME</u> | <u>BRMS</u> | <u>% in ROPE</u> | <u>Rhat</u> | <u>ESS</u> |
| Intensity | **(Intercept)** | 7.98 | 8.00 | 7.98 | [7.25, 8.72] | [7.25, 8.74] | [ 7.322, 8.618] | <0.001 | 0 | 100.00% | 0 | 1.001 | 1957.107 |
|  | Group | -0.81 | -0.85 | -0.71 | [-2.56, 0.93] | [-2.73, 1.02] | [-2.119, 0.745] | 0.361 | 0.354 | 78.18% | 21.992 | 1 | 2608.068 |
|  | **Cue** | 2.58 | 2.59 | 2.55 | [1.96, 3.20] | [1.96, 3.21] | [ 2.036, 3.054] | <0.001 | 0 | 100.00% | 0 | 1 | 6480.134 |
|  | *Trial* | -0.44 | -0.42 | -0.44 | [-0.54, -0.33] | [-0.52, -0.32] | [-0.526, -0.350] | <0.001 | 0 | 100.00% | 5.917 | 1 | 12483.224 |
|  | *Group \* Cue* | 1.28 | 1.27 | 1.20 | [-0.18, 2.75] | [-0.23, 2.77] | [ 0.055, 2.368] | 0.086 | 0.0958 | 94.65% | 10.083 | 1.001 | 6599.099 |
|  | Group * Trial | -0.15 | -0.19 | -0.15 | [-0.41, 0.10] | [-0.42, 0.04] | [-0.354, 0.059] | 0.232 | 0.1044 | 88.33% | 93.642 | 1 | 14314.655 |
|  | Cue * Trial | -0.03 | -0.08 | -0.03 | [-0.24, 0.18] | [-0.25, 0.09] | [-0.194, 0.153] | 0.791 | 0.3663 | 60.02% | 99.883 | 1 | 13010.665 |
|  | (Group * Cue) * Trial | 0.12 | 0.05 | 0.12 | [-0.39, 0.62] | [-0.35, 0.46] | [-0.275, 0.538] | 0.649 | 0.793 | 67.13% | 78.558 | 1 | 13453.963 |
| Unpleasantness | **(Intercept)** | 5.42 | 5.43 | 5.40 | [4.66, 6.18] | [4.67, 6.20] | [ 4.756, 6.055] | <0.001 | 0 | 100.00% | 0 | 1.001 | 2307.889 |
|  | Group | -0.09 | -0.12 | -0.10 | [-1.90, 1.72] | [-2.05, 1.81] | [-1.507, 1.350] | 0.924 | 0.9006 | 54.48% | 25.392 | 1.001 | 3741.232 |
|  | **Cue** | 1.48 | 1.47 | 1.46 | [1.13, 1.83] | [1.13, 1.80] | [ 1.140, 1.771] | <0.001 | 0 | 100.00% | 0 | 1 | 11086.081 |
|  | *Trial* | -0.32 | -0.31 | -0.32 | [-0.40, -0.24] | [-0.39, -0.23] | [-0.388, -0.252] | <0.001 | 0 | 100.00% | 20.75 | 1 | 16085.791 |
|  | **Group \* Cue** | 1.59 | 1.58 | 1.54 | [0.75, 2.42] | [0.78, 2.39] | [ 0.782, 2.278] | <0.001 | 0.0001 | 99.88% | 0.5 | 1 | 10754.967 |
|  | Group * Trial | -0.07 | -0.07 | -0.07 | [-0.27, 0.13] | [-0.26, 0.12] | [-0.233, 0.089] | 0.491 | 0.4627 | 75.87% | 98.308 | 1 | 16393.391 |
|  | Cue * Trial | -0.04 | -0.07 | -0.04 | [-0.21, 0.13] | [-0.21, 0.08] | [-0.176, 0.095] | 0.649 | 0.3664 | 68.16% | 99.792 | 1 | 15911.796 |
|  | (Group * Cue) * Trial | 0.05 | 0.03 | 0.05 | [-0.35, 0.45] | [-0.31, 0.36] | [-0.264, 0.380] | 0.8 | 0.8763 | 60.16% | 83.25 | 1 | 16260.922 |
| Skin<br>conductance<br>response<br>(CDA<br>Ampsum, Z-scored) | (Intercept) | -0.216 | -0.215 | -0.216 | [-0.29,<br>-0.14] | [-0.30,<br>-0.13] | [-0.278,<br>-0.152] | 0.000 | 0 | 100.00% | 0.033 | 1 | 10232.784 |
|  | Group | -0.021 | -0.021 | -0.021 | [-0.20,<br>0.15] | [-0.24,<br>0.20] | [-0.165,<br>0.129] | 0.821 | 0.8401 | 59.20% | 58.458 | 1 | 9689.083 |
|  | Cue | 0.132 | 0.137 | 0.132 | [0.00,<br>0.26] | [0.03,<br>0.24] | [0.028,<br>0.242] | 0.047 | 0.0109 | 97.62% | 20.467 | 1 | 17008.287 |
|  | Trial | -0.054 | -0.058 | -0.054 | [-0.08,<br>-0.02] | [-0.09,<br>-0.03] | [-0.079,<br>-0.029] | 0.001 | 0.0001 | 99.98% | 92.767 | 1 | 17267.529 |
|  | Group * Cue | 0.027 | -0.002 | 0.026 | [-0.28,<br>0.34] | [-0.25,<br>0.25] | [-0.233,<br>0.267] | 0.863 | 0.9864 | 56.73% | 36.283 | 1 | 14843.577 |
|  | Group *<br>Trial | -0.031 | -0.033 | -0.031 | [-0.10,<br>0.04] | [-0.10,<br>0.04] | [-0.092,<br>0.030] | 0.408 | 0.3476 | 79.61% | 88.325 | 1 | 16533.377 |
|  | Cue * Trial | -0.024 | -0.027 | -0.023 | [-0.09,<br>0.04] | [-0.08,<br>0.03] | [-0.073,<br>0.029] | 0.449 | 0.3093 | 77.38% | 95.1 | 1 | 15037.218 |
|  | (Group *<br>Cue) * Trial | -0.059 | -0.068 | -0.059 | [-0.21,<br>0.09] | [-0.19,<br>0.06] | [-0.179,<br>0.058] | 0.432 | 0.2862 | 78.56% | 56.292 | 1 | 14678.671 |

<sup>f</sup>. This table presents results of linear mixed models predicting subjective pain (intensity or unpleasantness) and skin conductance response (Z-scored summary of amplitudes) as a function of Cue (High vs Low), Group (VMPFC vs Healthy Control), and Trial. Separate models were conducted for each outcome measure. See Table 2 for additional information on analytic approaches.

There was no main effect of Group in any model (all p’s > 0.3), indicating that the impact of VMPFC lesions was specific to expectations. None of the remaining interaction terms were significant (all p’s > 0.1; see Table 3). Thus, contrary to our second hypothesis and consistent with group differences in self-reported expectations, the VMPFC group reported significantly *greater* cue effects on pain unpleasantness (see Figure 4).

**Figure 4.**
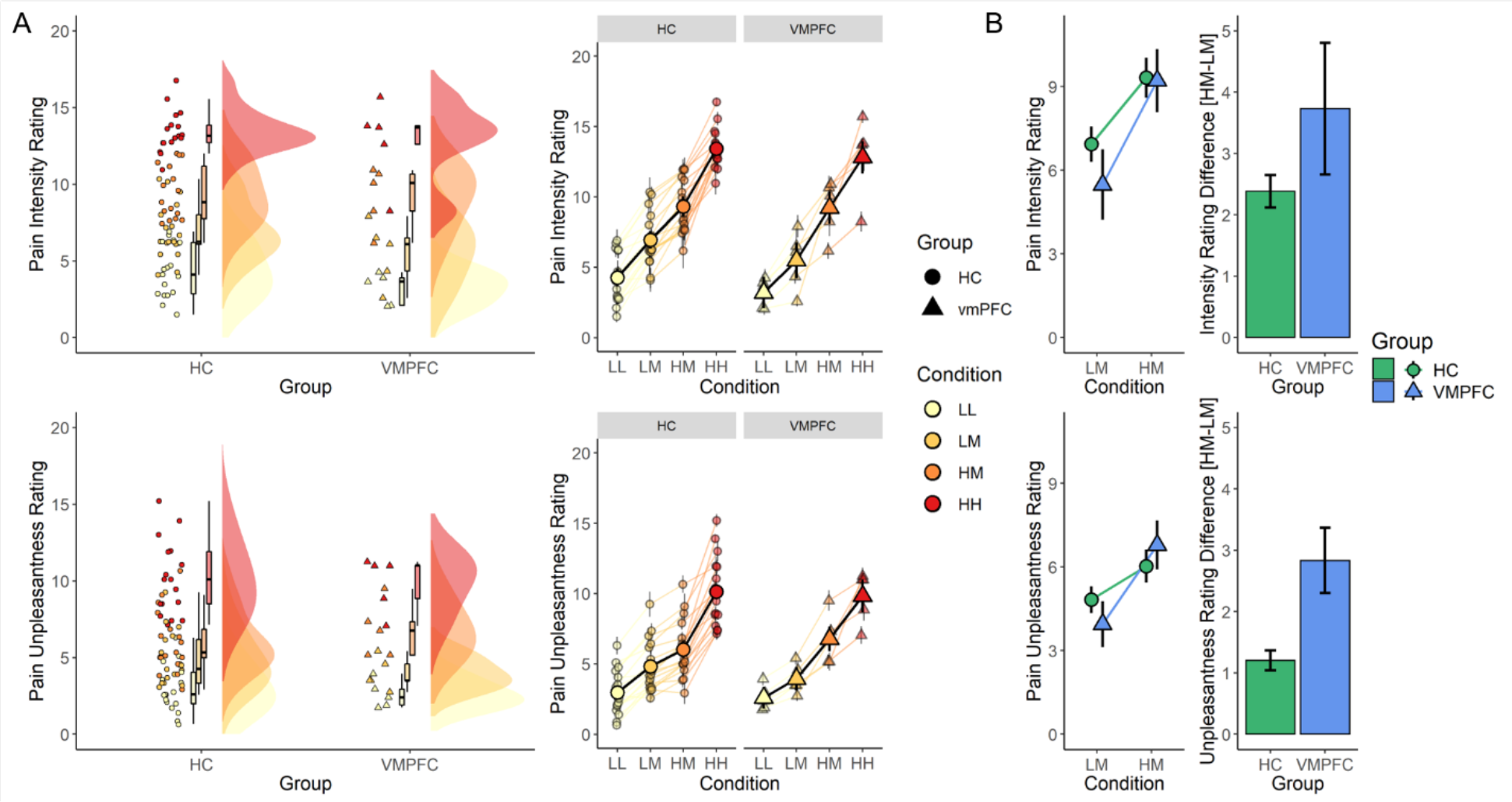
Behavioral Results during the expectancy task. **(A)** Individual subject data. Modified raincloud plots (left) showing the distribution of pain intensity (top) and unpleasantness (bottom) ratings across all 4 conditions (LL, LM, HM, and HH conditions represented from yellow to red). Line graph of pain ratings by condition (right) showing group mean ± s.e.m. ratings superimposed on individual subject values. Note the consistency of ratings across individual subjects and greater separation between LM (light orange) and HM (dark orange) peaks in the VMPFC group relative to the HC group. **(B)** Group summary data. Line (right) and bar (left) graphs showing differences between ratings for the critical medium heat trials preceded by a low (LM) or high (HM) pain cue, showing greater expectancy effect on unpleasantness ratings in the VMPFC group (blue triangles) relative to the HC group (green circles). Similar findings were observed for intensity ratings, but did not meet frequentist criteria for statistical significance. Statistics are reported in Table 5.

### Individuals with VMPFC lesions display blunted skin conductance responses overall but similar autonomic sensitivity to heat intensity and expectancy

As a third test of Hypothesis 2, we evaluated whether groups differed in the extent to which autonomic responses to thermal stimuli are modulated by pain-predictive cues. In addition, to formally test our third hypothesis (i.e., that VMPFC lesions would impact the association between pain and autonomic responses), we examined autonomic responses as a function of temperature across all trials, and as a function of subjective pain ratings within medium heat trials.

We first measured autonomic responses as a function of temperature across all trials as a basis for subsequent group comparisons. Whereas we observed practically significant effects of temperature on SCR amplitude regardless of statistical approach (all p’s < 0.001; see Supplementary Table S3), there was no influence of temperature on heat-evoked mean HR change (see Supplementary Figure 1). We therefore focused on SCR for the remainder of our autonomic analyses. Analyses of square-root normalized SCR amplitude revealed group differences in the intercept that were practically significant, indicating that SCR was blunted in the VMPFC lesion group (see Supplementary Table S3), consistent with prior reports^33, 35^. We therefore z-scored responses within participants to account for group differences in mean SCR amplitude in subsequent analyses. Importantly, there were no group differences in the magnitude of temperature effects on SCR when we analyzed z-scored SCR amplitudes (all p’s > 0.1; see Figure 5 and Supplementary Table S3). For additional results across temperature and to compare SCR amplitude with other phasic SCR measures, see Supplementary results.

**Figure 5.**
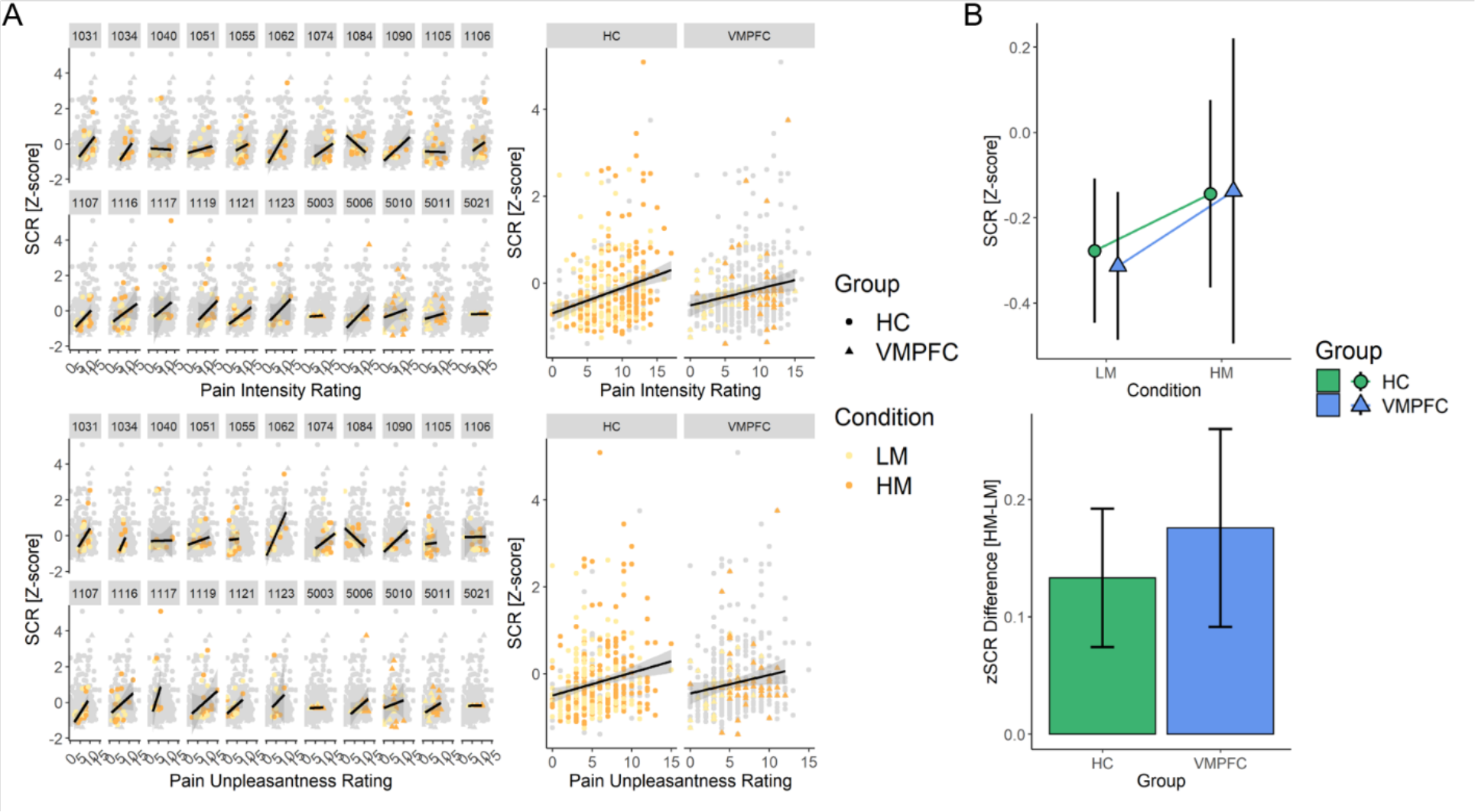
Autonomic Results during the expectation task. **(A)** Individual subject scatter plots (left) between intensity (top) and unpleasantness (bottom) ratings and z-scored SCR, and group summary scatter plots (right) showing positive relationship between pain ratings and SCR amplitudes for the subset of medium temperature trials in both groups. LM trials are shown in yellow and HM trials shown in orange. **(B)** Group summary plots showing greater SCR amplitudes for medium trials preceded by high (HM) relative to low (LM) pain cues (top) in both groups and no differences between groups in the difference in SCR amplitude between HM and LM trials (bottom).

Next, we explored whether VMPFC lesions impacted expectancy effects on autonomic responses, consistent with effects on subjective pain ratings. Analyses of SCR amplitude on medium heat trials revealed a main effect of Cue (all p’s < 0.05; see Table 3), such that SCR amplitude was higher in response to medium heat paired with high pain cues relative to low pain cues, although this effect was of undecided practical significance based on Bayesian posterior estimates (20.47% in ROPE). Interestingly, there were no differences between Groups, nor any interactions with Group (all p’s > 0.2), suggesting that groups did not differ in the magnitude of cue effects on SCR (Figure 5), although they did differ in cue effects on pain. Analyses of square-root normalized data are reported in Supplementary Table S4 and are consistent with findings from z-scored outcomes, with the addition of group differences in the intercept; however, these can be accounted for by overall blunting of the SCR in the VMPFC group and we therefore do not use them for inference. We observed consistent findings for other phasic SCR measures, which each indicated significant differences as a function of Cue and no influence of Group (see Supplementary Table S4). Thus, across outcome measures, we observed larger heat-evoked SCRs when moderate heat was preceded by high pain cues relative to low pain cues.

Surprisingly, after accounting for the overall reduction in SCR amplitude, the VMPFC group exhibited similar cue effects on SCR relative to the HC group, indicating that a failure to generate autonomic responses did not account for the observed differences in pain ratings.

Finally, to directly test our third hypothesis, we evaluated whether the relationship between SCR and subjective pain ratings differed between groups. We focused on responses to medium heat trials. When we z-scored responses to account for group differences in mean SCR amplitude, we observed significant positive associations with each rating for all SCR outcomes (all p’s < 0.1; see Supplementary Table S5) but no group differences in the association between SCR amplitude and pain intensity or unpleasantness ratings (all p’s > 0.3). As above, we did observe group differences when we used square root normalized SCR values, but these reflect overall blunting in the VMPFC group (see Figure 5 and Supplementary Table S5). Thus, in contrast to our third hypothesis, VMPFC lesions were not associated with differences in the correspondence between subjective pain and physiologic arousal.

## Discussion

Using a combination of subjective pain ratings and physiologic measures of autonomic reactivity in a unique population of neurosurgical patients with focal, bilateral damage to VMPFC, we present evidence supporting a critical role of VMPFC in modulating and updating conscious expectancies about pain. Our study provides the first evidence that VMPFC lesions: 1) enhance explicit expectations about painful stimuli, and 2) directly influence the magnitude of expectation effects on the conscious perception of such stimuli. For an identical level of thermal stimulation, pain ratings were driven more strongly by the expected level of pain in patients with VMPFC lesions than in neurologically healthy adults. These differences were particularly strong for pain unpleasantness, which reflects the affective and motivational qualities of the pain experience. Patients with VMPFC lesions reported larger differences in explicit expectations based on verbal instructions alone, and these differences remained stable over the course of the experiment. Notably, differences in expectations about pain were not linked to altered pain sensitivity overall, nor to altered modulation of autonomic responses. While physiological arousal was blunted overall in patients with focal VMPFC lesions and pain tolerance was somewhat (though not significantly) higher, general nociceptive processing was unaffected when these subtle differences were considered. Here, we discuss these findings and their implications.

### VMPFC is involved in generating and updating expectations about pain

The specific mechanisms through which expectations shape pain perception has been the focus of considerable prior research. FMRI and PET studies indicate that expectations based on pain-predictive cues or placebo administration shape responses to noxious stimuli in regions within the so-called pain matrix, including the insula, thalamus, and dorsal anterior cingulate cortex (dACC)^12^. Our previous work suggests that cue effects on these regions predict subjective pain ratings and, importantly, are mediated by cue responses in the VMPFC, as well as the striatum^17^. Other studies have also linked expectations and contextual modulation of pain to the VMPFC (for review, see ^31, 59^). Some of these studies highlight connectivity between VMPFC or adjacent rostral ACC and midbrain periaqueductal gray (PAG) as a predictor of individual differences in the strength of the expectancy effect, suggesting that expectations of pain or relief may in part reflect differential recruitment of known brainstem pain control circuits that directly impinge on nociceptive transmission at the level of the dorsal horn^17, 60–63^. Here we show that, while lesions of the VMPFC have a causal influence on expectancy effects in pain, the result is one of *enhancement* rather than attenuation, and that this impacts both expectations for pain exacerbation and pain relief, rather than being strictly linked to antinociceptive circuits. Our findings suggest that a core function of VMPFC is to modulate and update expectations based on integrating context and prior experience with somatosensory information^31^, particularly in the context of judgements about pain unpleasantness.

A primary role of VMPFC in updating expectancies is consistent with computational models of reinforcement learning that implicate VMPFC subregions, particularly OFC, in processing expected value and learning latent rules^23, 24, 64^. Our results provide new evidence that the VMPFC not only shapes behavior (e.g., choices for valued options), but that it also shapes explicit, subjective predictions about valued outcomes. Participants with VMPFC lesions reported larger differences in expectations as a function of cue type both immediately after instructions (prior to learning) and throughout the test phase of the experiment (i.e., when high and low cues were intermittently paired with medium intensity heat). Thus, our findings suggest that VMPFC’s causal role is not only in generating *latent* predictions, but also in updating explicit expectations based on instructions and learning. These findings are largely consistent with previous views of the OFC/VMPFC and its role in generating and updating expected value and abstract rules, but highlight that acquisition of explicit (as opposed to latent) expectations does not require intact VMPFC.

In contrast to our hypothesis that lesions to the VMPFC would *reduce* the extent to which expectations would shape aversive outcomes, we found that VMPFC lesions *enhanced* cue-based expectations and expectancy-based modulation. Unlike previous work on the role of VMPFC in associative learning, participants in our task receive instructions *and* experiential learning. It is possible that the effects of instructions were mediated by intact regions such as the dorsolateral prefrontal cortex (DLPFC), which has been implicated in instructed learning,^60, 65, 66^ cognitive control^67^, and expectancy-based pain modulation^17, 68, 69^, and that the VMPFC lesions impact the extent to which actual reinforcement *reduces* the effects of verbal instructions. This would account for the observation that group differences in the test phase appear to be driven by a reduction in expectancy effects among neurologically healthy comparison adults immediately following the introduction of moderate heat trials, potentially reflecting the successful integration of new sensory feedback to update the previously instructed cue-related expectancies, whereas this learning is not observed in the VMPFC lesion group. Unfortunately, our task was not designed to dissociate the effects of verbal instructions from experiential learning, and therefore we cannot determine whether VMPFC patients relied more strongly on instructions or associative learning. We previously found that BOLD responses in the OFC tracked expected value that updated from both learning and instruction,^60^ indicating that both sources of expectation can impact responses in this region. Future work could incorporate instructed reversals^70–72^ or compare instructed *vs.* learned expectancies to separately isolate regional effects on instruction from effects on experience.

The fundamental role of the VMPFC in linking expectations with affectively salient outcomes is consistent with a large body of lesion work in primate and rodent models^73–76^, as well as human neuroimaging studies^77–79^. Though the vast majority of prior work linking VMPFC lesions with deficits in prediction and valuation focus on appetitive processes, including reward-based learning and responses to positive hedonic outcomes, human imaging work implicates the same region in predicting aversive outcomes ^17, 60, 80, 81^. Studies that combine appetitive and aversive learning indicate that the OFC contains neurons that respond to both types of outcomes and signal outcome valence^82, 83^. We hypothesize that our findings are unlikely to be specific to pain, and that lesions of this region are likely to have similar effects on expectancy-based modulation of appetitive stimuli, such as hedonic tastes or other rewarding outcomes. This should be directly tested in future empirical work.

### VMPFC lesion effects are most pronounced for judgements about the affective qualities of pain

Most work in quantitative sensory testing has focused on evaluating pain intensity^84–86^. Here, we modified an adaptive calibration QST procedure^17, 43, 47, 48^ and trained participants to separately evaluate sensory and affective components of pain (i.e., pain intensity and unpleasantness, respectively), which allowed us to specifically test for altered affective processing in VMPFC lesion patients. Although the sensory and affective components of pain are highly correlated in most situations, there is a substantial body of evidence indicating that they are dissociable. Separate brain regions track intensity and unpleasantness ratings during acute pain based on fMRI and PET studies^5, 87, 88^, and preclinical models identify specific circuits in the amygdala and rostral anterior cingulate that selectively modulate pain affect while leaving sensory processing intact^89–91^. Importantly, historical reports of patients with frontal lobotomy indicate that frontal lesions can drastically alter the affective components of pain without impacting sensory/discriminative judgements about stimulus location and intensity^2^. Consistent with its proposed role in generating pain affect, we found that the impact of VMPFC lesions on cue-based expectations was most pronounced for ratings of pain unpleasantness. However, we did not observe more general deficits in pain affect as might be expected from the prior clinical and fMRI literature, suggesting that the causal role of VMPFC is likely specific to the link between expected value and affective processing, rather than simply mediating subjective judgements about unpleasantness, per se.

Unexpectedly, we found no evidence of widespread alterations in pain processing in our sample of patients with focal VMPFC lesions. Although pain thresholds were somewhat elevated in the VMPFC lesion group, there were no significant differences between groups in pain thresholds or tolerance, and temperature-pain associations were similar between individuals with and without VMPFC lesions. This is contrary to findings from prior animal studies suggesting that inactivating regions homologous to VMPFC and the adjacent ACC in rodents elicits significant reductions in pain avoidance behaviors^90, 92, 93^, and with historical reports of human neurosurgical patients noting profound asymbolia for pain following frontal lobotomy. We see three possible explanations for this apparent discrepancy. The first relates to neuroanatomy. As we note elsewhere, our VMPFC sample is unique in terms of the focality and homogeneity of lesions; all subjects had bilateral VMPFC damage from surgical resection of orbital meningiomas, leaving adjacent structures like DLPFC (described above) and rACC largely intact. VMPFC and rACC are often conflated in the literature, which is further compounded by issues with regional homology across species^94–96^. Notably, in all prior human neurosurgical cases, lesions of ACC and VMPFC, ACC alone, or the cingulum bundle connecting ACC to more posterior brain regions were necessary for the observed reductions in pain affect^1, 2, 8–10^.

Further, in both humans and animal models, individual neurons within the ACC (but not VMPFC or rodent homologue infralimbic cortex) have been shown to encode pain^90, 97^. The absence of more profound effects on pain processing and pain thresholds suggests that spared regions of ACC in our sample are more likely than VMPFC to mediate these effects. The second and third potential explanations relate to the testing equipment and experimental design, respectively. For safety purposes, the Medoc software includes a maximum temperature limit of 49° C at the stimulus duration we used, which was lower than the computed subjective pain threshold for three of the five VMPFC lesion patients. Therefore, the computed pain threshold may underestimate the true threshold for VMPFC lesion patients. Finally, three HC participants with an r^2^ > 0.4 at the end of the calibration procedure were included in the calibration analyses but excluded from the task analyses to maintain consistency with prior work^17, 43, 98^. When we exclude these participants, we do find a main effect of group (p=0.046), such that VMPFC lesion participants have elevated pain tolerance, suggesting that “noisy” calibration values among excluded HC participants may be artificially suppressing a true group difference in pain processing in patients with VMPFC lesions. However, post-hoc tests at each temperature level remain non-significant even after removing the three HC participants, suggesting that their overall contribution is modest. Future studies involving patients with lesions involving multiple subregions of PFC, higher intensities of pain stimulation, or different pain modalities are needed to more conclusively determine whether focal VMPFC lesions increase pain thresholds.

### Patients with VMPFC lesions have attenuated autonomic responses to heat but show intact associations with pain and modulation by expectancy

One proposed mechanism by which VMPFC lesions may impact pain perception is through impairment of the integration of emotion, conceptualized by Damasio as the somatic and visceral ‘body state’, with conscious expectations about pain and incoming sensory information to guide subjective experience and behavior^33^. Damasio and others have postulated that VMPFC plays a key role in linking peripheral physiological components of an emotional response (so-called “somatic markers”) with higher-order cognitive processes to guide decision-making and behavior^31, 99^. Previous work indicates that patients with VMPFC lesions fail to mount appropriate skin conductance responses as uncertain choices evolve to become more risky and less rewarding, which corresponds with a failure to update their decision-making strategy toward more favorable outcomes^34, 36^. We found that heat-evoked skin conductance responses were indeed blunted in the VMPFC lesion group. However, once differences in mean amplitude were accounted for, both groups showed similar effects of temperature and pain-predictive cues, and no differences in the correspondence between autonomic responses and subjective decisions (i.e., pain report). These findings diverge from previous findings of expectancy driven effects on physiological responses published in our lab^100^, and indicate that differences in autonomic responses cannot account for the differences in expectations or expectancy effects on subjective pain. Future work should explore whether the differences in these findings may be explained by the inherently increased averseness of pain as compared to other aversive stimuli (e.g., emotional pictures), or by specific neuroanatomical substrates that were not damaged in our sample.

## Limitations and future directions

One feature of this study that warrants consideration is the limited sample size of VMPFC lesion patients (n = 5). We used extremely stringent selection criteria for our target group; lesions had to involve substantial portions of the VMPFC but could not extend significantly outside the VMPFC and could not involve other brain regions implicated in pain processing. Therefore, although our sample size may be small by conventional VMPFC lesion patient standards, which typically feature n = 5 to n =12 VMPFC lesion patients, it is unique with respect to the homogeneity, uniformity, and focality of VMPFC lesions. In addition, all lesions were bilateral, in contrast to other studies that include only lateralized lesions^8^. Future work in larger samples of VMPFC lesion patients with voxel-based lesion-symptom mapping^101^ (VLSM) is needed to more conclusively determine the relationship between damage to specific subregions of VMPFC and individual components of expectancy and pain.

Taken together, these results highlight a critical role of VMPFC in the processing of affective responses to painful stimuli. Our unexpected finding of an increase in the strength of effect of pain-predictive cues on expectations and pain in the VMPFC lesion group suggests that these patients relied more heavily on instructions and conscious expectation than incoming sensory or nociceptive information in selecting their ratings of painful stimuli. Although contrary to our initial hypothesis, this finding is remarkably consistent with a substantial learning, reward, and valuation literature that proposes a central role for VMPFC in integrating sensory information with contextual information to guide action tendencies. Our findings support a key role of VMPFC in modulating and updating expectations in aversive contexts and highlight that other processes commonly ascribed to VMPFC, including autonomic integration and encoding of pain affect, likely rely on neural substrates outside of the region of damage in our sample.

## Supporting information

Supplemental Methods, Results, Tables, and Figures

## Acknowledgements

The authors would like to thank Maia Pujara for assistance with data collection, Joe Weilgosz for assistance with technical implementation, and Geoff Schoenbaum, Allan Basbaum, and Howard Fields for their feedback on a prior version of this manuscript.

## Abbreviations

HH: High Pain Cue + High Heat
HM: High Pain Cue + Moderate Heat
LM: Low Pain Cue + Moderate Heat
LL: Low Pain Cue + Low Heat
VMPFC: Ventromedial prefrontal cortex;

