## Supplemental Methods, Results, Tables, and Figures for "Human ventromedial prefrontal cortex lesions enhance expectation-related pain modulation"

**Supplementary Materials to accompany "Human ventromedial prefrontal cortex lesions enhance expectation-related pain modulation**

Motzkin JC[1,2], Hiser J[3], Carroll I[4], Wolf R, Baskaya MK[5], & Koenigs M[‡3] & Atlas LY[6,7,8‡*],

#### Supplementary Methods

**Multilevel linear model specification.** Linear mixed models were consistent across analytic approaches (LMER[1], NLME[2], and BRMS[3]) and across outcome measures, with few exceptions. All models included fixed effects of Group with Subject modeled as random. Effects of time (Trial or Block) were modeled as fixed in all models, while we included random slopes for other factors unless models were unable to converge.

Analyses that included all trials across temperature had identical specifications whether evaluated by LMER, NLME, or BRMS. All included random slopes for effects of Temperature on outcomes (intensity, unpleasantness, or SCR). We also included a random effect of Cue in analyses across of pain ratings across all trials. Due to issues with convergence, we did not include a fixed effect of Trial when analyzing data from the calibration task, and did not include an effect of Cue when analyzing temperature effects on SCR.

All analyses restricted to medium heat trials all included random slopes for effects of Cue on pain outcomes (intensity, unpleasantness), with the exception of LMER analyses of unpleasantness on medium heat trials which would not converge unless random slopes were omitted. We included only random intercepts for all analyses of SCR outcomes on medium heat trials due to issues with convergence.

Confidence intervals for LMER results were acquired using the "tab_model" function from the R package "sjPlot" [4]. Confidence intervals for NLME results were obtained using the 'intervals' function from the package "nlme" [2]. Results of BRMS models were obtained using bayestestR[5] to evaluate posteriors and evaluate practical significance. As discussed in

Makowski *et al.*[5,6], the probability of direction is comparable to a frequentist p-value to evaluate statistical significance; we therefore provide this value in all tables for comparison with frequentist approaches (i.e. LMER and NLME mixed models). “Practical significance” can be evaluated using the region of partial equivalence, or ROPE, which is defined as a range around a negligible parameter value (in our case, 0) that depends on the standard deviation of the outcome “y” (ROPE = [-0.1*$SD_y$; 0.1*$SD_y$]). We evaluated the percent of posterior estimates falling within the full ROPE range, and therefore defined practical significance (i.e. ability to reject the null hypothesis of no effect) as fewer than 2.5% of posterior estimates falling within the ROPE, and would accept the null if >97.5% of estimates fell within the ROPE[5,6].

#### Supplementary Results

**Expectancy rating analysis.**

In addition to analyses of difference scores, we also analyzed expectancy ratings as a function of cue during the test phase (i.e. blocks 3-8, including medium heat trials) using ANOVAs and linear mixed models. ANOVAs revealed a significant main effect of Cue on both expected intensity ($F(1,20) = 98.218$, $p < .001$) and expected unpleasantness ($F(1,20) = 108.166$, $p < .001$). There was a significant Group x Cue interaction in expected unpleasantness ($F(1,20) = 8.475$, $p = .008$) and a marginal Group x Cue interaction in expected intensity ($F(1,20) = 4.149$, $p = .055$). Post-hoc pair-wise comparisons for both intensity and unpleasantness indicated that each group reported significant differences in expectations as a function of cue, both for intensity ratings (VMPFC high cue expectancy - VMPFC low cue expectancy: $B = 10.56$, $t = 4.1$, $p = .001$; Control Group high cue expectancy - Control Group low cue expectancy: $B = 6.73$, $t = 4.82$, $p < .001$) and for unpleasantness ratings (VMPFC high cue expectancy - VMPFC low cue expectancy: $B = 7.37$, $t = 3.24$, $p = .014$; Control Group high cue expectancy - Control Group low cue expectancy: $B = 4.97$, $t = 4.03$, $p = .002$). There were no Group differences in response to either cue alone (all p’s > 0.5), nor any Group x Cue interactions (all p’s > 0.05). Thus groups differed primarily in the magnitude of the differential response, rather than specifically to high or low pain cues.

Linear mixed models revealed similar results (see Supplementary Table S2). Relative to low pain cues, high pain cues were associated with greater expected intensity and unpleasantness and this effect was practically significant (Main effect of Cue: all p’s < .001; 0% in ROPE). As shown in Figure 4, individuals with VMPFC lesions reported stronger differences in expected unpleasantness as a function of cue and this effect was practically significant (Group x Cue: all p’s < .001; 2.28% in ROPE). Group differences in cue effects on expected intensity ratings ranged from marginal to significant depending on the linear mixed model approach (see Supplementary Table S2) and were of undecided significance based on Bayesian models (8.61% in ROPE). There were no main effects of Group or Block in either model (all p’s > 0.1).

**Analyses of temperature effects on SCR.**

The main manuscript focuses on temperature effects on square-root normalized and Z-scored SCR amplitude, as computed by the sum of amplitudes from Ledalab’s Continuous Decomposition Analysis (CDA[7]). We also evaluated temperature effects on other phasic SCR measures derived from CDA, including SCR and tonic mean, as well as the sum of amplitudes derived from trough-to-peak scoring in Ledalab. We focus on measures that were z-scored to account for differences in the average amplitude, as reported in the main manuscript. As reported in Supplementary Table S3, we observed significant associations between heat-evoked SCR and temperature in all models (all p’s < .001), and temperature effects did not differ by Group for any SCR measure.

**Analyses of non-normalized SCR outcome measures in response to medium heat trials.**

We measured whether groups differed in the magnitude of expectancy effects on heat-evoked SCR by focusing on cue effects during the critical medium heat trials. Our main manuscript focuses on z-scored outcome measures to account for SCR blunting in individuals with VMPFC lesions, whereas here we report results on non-normalized data. We still square-root normalized all outcomes to account for non-normality, however we did not z-score results to retain group differences in overall amplitude.

We first tested the sum of CDA amplitude estimates. Consistent with results across temperatures, we observed a main effect of Group ($\beta$=-0.48, SE=0.16; p = .008), such that SCR amplitudes were higher in control participants relative to participants with VMPFC lesions. Consistent with z-scored results, we observed a main effect of Cue ($\beta$=0.065, SE=0.033; p = .049), such that amplitudes were higher in response to high pain cues relative to low pain cues, and a main effect of Trial ($\beta$=-0.025, SE=0.008; p = .001), such that SCR amplitude decreased over the course of the task. There were no interactions between Group and cue, trial, or three-way interactions (all p's > 0.5). We observed similar results when we tested square-root normalized SCR and when we tested the sum of amplitude estimates using trough-to-peak estimates (see Supplementary Table S4). We also tested cue effects on tonic mean as estimated by CDA. In contrast to phasic SCR measures, we did not observe significant cue effects on tonic SCR or any differences by Group (see Supplementary Table S4). The only predictor that significantly modulated tonic SCR was Trial ($\beta$=-0.002, SE=0.000; p < .001, such that tonic SCR decreased over time.

### Supplementary Figures

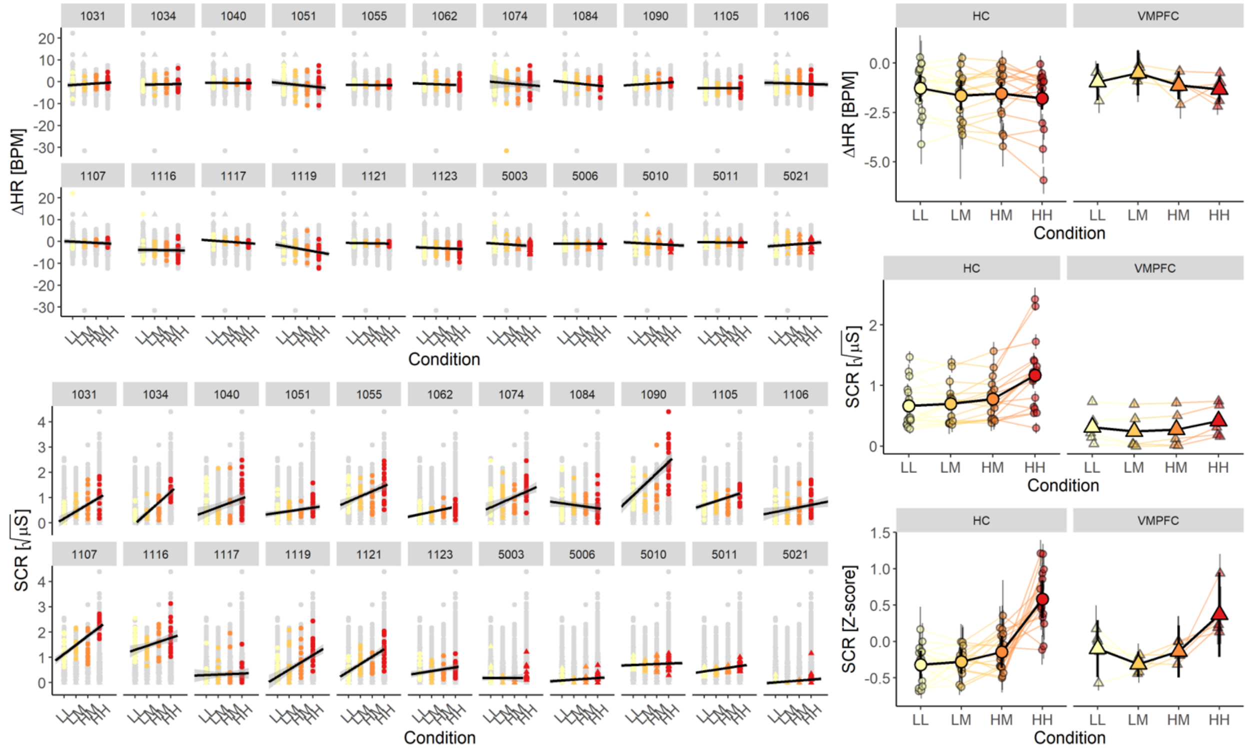


**Supplementary Figure 1. Autonomic Responses to thermal pain stimuli.** *Individual subject scatter plots (left) between stimulus conditions and trial-wise change in HR (top) and square root transformed SCR (bottom). Group summary line graphs (right) depicting autonomic responses by condition for change in HR (top), square-root transformed SCR (middle), and z-scored SCR (bottom) in each group. There is no relationship between HR and temperature, no differences between groups, so HR was excluded from subsequent analyses. Root transformed SCR maintains group differences in response amplitude, but patterns of responses are more similar between groups when responses are normalized within groups. SCR=skin conductance response, μS=microsiemens, ΔHR=change in heart rate, BPM=beats per minute, LL=Low Pain Cue+Low Heat, LM=Low Pain Cue+ Moderate Heat, HM=High Pain Cue +Moderate Heat. HH=High Pain Cue + High Heat.*

#### Supplementary Tables

Supplementary Table S1. Temperature effects on pain during the adaptive calibration.[a]

| | | Estimates | | | Confidence Intervals | | | P-values / Probability of direction | | | Bayesian estimates | | |
|---|---|---|---|---|---|---|---|---|---|---|---|---|---|
| Outcome measure | Effect | LMER | NLME | BRMS | LMER | NLME | BRMS | LMER | NLME | BRMS | % in ROPE | Rhat | ESS |
| Intensity | **(Intercept)** | 8.475 | 8.428 | 8.49 | 7.17 – 9.78 | [7.08, 9.78] | [ 7.18, 9.90] | 0.000 | 0.000 | 100% | 0% | 1.001 | 2277 |
| | Group | -1.113 | -1.115 | -0.99 | -2.75 – 0.52 | [-2.90, 0.67] | [-2.58, 0.72] | 0.196 | 0.208 | 88.15% | 21.48% | 1 | 3487 |
| | **Temperature** | 1.165 | 1.242 | 1.16 | 1.02 – 1.32 | [1.08, 1.40] | [ 1.00, 1.33] | 0.000 | 0.000 | 100% | 0% | 1 | 5609 |
| | Group x Temperature | 0.078 | 0.067 | 0.07 | -0.11 – 0.27 | [-0.14, 0.27] | [-0.14, 0.27] | 0.437 | 0.521 | 76.44% | 99.98% | 1.001 | 7078 |
| Unpleasantness | **(Intercept)** | 5.507 | 5.447 | 5.53 | 4.45 – 6.57 | [4.37, 6.52] | [ 4.34, 6.62] | 0.000 | 0.000 | 100% | 0% | 1.002 | 2328 |
| | Group | -0.881 | -0.887 | -0.81 | -2.21 – 0.45 | [-2.31, 0.53] | [-2.16, 0.57] | 0.207 | 0.210 | 88.19% | 21.22% | 1 | 3397 |
| | **Temperature** | 0.842 | 0.872 | 0.84 | 0.71 – 0.97 | [0.74, 1.01] | [ 0.69, 0.98] | 0.000 | 0.000 | 100% | 0% | 1.001 | 5904 |
| | Group x Temperature | 0.072 | 0.080 | 0.07 | -0.10 – 0.24 | [-.09, 0.25] | [-0.12, 0.26] | 0.412 | 0.364 | 77.88% | 99.85% | 1 | 6656 |

[a]. This table presents results of linear mixed models predicting subjective pain as a function of mean-centered Temperature and Group (VMPFC vs Healthy Control) based on ratings during the Adaptive Staircase Calibration task prior to the main experiment. Separate models were conducted using Intensity ratings and Unpleasantness ratings.

Supplementary Table S2. Explicit expectancy during test phase (blocks 3-8).[b]

| | | Estimates | | | Confidence Intervals | | | P-values / Probability of direction | | | Bayesian estimates | | |
|---|---|---|---|---|---|---|---|---|---|---|---|---|---|
| Outcome measure | Predictors | LMER | NLME | BRMS | LMER | NLME | BRMS | LMER | NLME | BRMS | % in ROPE | Rhat | ESS |
| Expected intensity | (Intercept) | 8.598 | 8.593 | 8.51 | 7.55 – 9.65 | [7.53, 9.65] | [ 7.31, 9.64] | 0.000 | 0.000 | 100% | 0% | 1.002 | 2532 |
| | Block | -0.018 | -0.013 | -0.02 | -0.14 – 0.10 | [-0.12, 0.09] | [-0.14, 0.10] | 0.764 | 0.804 | 61.99% | 100% | 1 | 13337 |
| | Cue | 7.242 | 7.231 | 6.66 | 5.89 – 8.59 | [5.87, 8.60] | [ 5.07, 8.09] | 0.000 | 0.000 | 100% | 0% | 1 | 4035 |
| | Group | -0.451 | -0.450 | -0.44 | -2.96 – 2.06 | [-3.13, 2.23] | [-2.84, 1.94] | 0.729 | 0.730 | 64.75% | 30.59% | 1.001 | 3999 |
| | Cue*Block | 0.065 | 0.082 | 0.06 | -0.17 – 0.30 | [-0.19, 0.35] | [-0.17, 0.30] | 0.589 | 0.548 | 70.19% | 100% | 1 | 13233 |
| | Group*Block | 0.079 | 0.073 | 0.08 | -0.20 – 0.36 | [-0.18, 0.32] | [-0.20, 0.37] | 0.585 | 0.562 | 70.93% | 99.78% | 1 | 14551 |
| | *Group*Cue* | 3.353 | 3.366 | 2.29 | 0.13 – 6.58 | [0.11, 6.62] | [-0.56, 5.15] | 0.055 | 0.043 | 93.47% | 8.61% | 1 | 5645 |
| | Group*Cue*Block | 0.086 | 0.082 | 0.09 | -0.48 – 0.65 | [-0.56, 0.72] | [-0.45, 0.66] | 0.764 | 0.801 | 61.97% | 90.09% | 1 | 13666 |
| Expected unpleasantness | (Intercept) | 6.621 | 6.619909 | 6.55 | 5.65 – 7.59 | [5.64, 7.59] | [ 5.51, 7.60] | 0.000 | 0.000 | 100% | 0% | 1.002 | 2276 |
| | Block | -0.077 | -0.075028 | -0.08 | -0.20 – 0.04 | [-0.19, 0.04] | [-0.19, 0.05] | 0.211 | 0.204 | 89.43% | 100% | 1 | 12773 |
| | Cue | 5.318 | 5.315257 | 5.1 | 4.33 – 6.31 | [4.32, 6.31] | [ 3.97, 6.17] | 0.000 | 0.000 | 100% | 0% | 1 | 4336 |
| | Group | 0.210 | 0.207941 | 0.09 | -2.10 – 2.52 | [-2.26, 2.67] | [-2.09, 2.37] | 0.861 | 0.862 | 53.33% | 29.12% | 1.002 | 3805 |

| | | | | | | | | | | | | | |
|---|---|---|---|---|---|---|---|---|---|---|---|---|---|
| | Cue*Block | 0.036 | 0.039808 | 0.04 | -0.20 – 0.28 | [-0.21, 0.29] | [-0.20, 0.28] | 0.766 | 0.753 | 61.91% | 99.85% | 1 | 11652 |
| | Group*Block | 0.058 | 0.055679 | 0.06 | -0.23 – 0.34 | [-0.22, 0.33] | [-0.22, 0.35] | 0.688 | 0.692 | 65.65% | 98.88% | 1 | 12316 |
| | **Group*Cue** | 3.514 | 3.518873 | 2.78 | 1.15 – 5.88 | [1.14, 5.90] | [ 0.46, 5.15] | 0.009 | 0.004 | 98.42% | 2.28% | 1.001 | 4659 |
| | Group*Cue*Block | -0.202 | -0.19743 | -0.2 | -0.77 – 0.37 | [-0.79, 0.40] | [-0.79, 0.37] | 0.488 | 0.514 | 74.99% | 73.69% | 1 | 12484 |

[b]. This table presents results of linear mixed models predicting subjective expectation rating as a function of Cue (High vs Low), Group (VMPFC vs Healthy Control), and Block (3-8, mean centered) based on ratings during the Adaptive Staircase Calibration task prior to the main experiment. Separate models were conducted using Intensity ratings and Unpleasantness ratings.

Supplementary Table S3. Temperature effects on SCR.[c]

| Outcome measure | | CDA: Ampsum (square root normalized) | | | CDA: SCR (square root normalized) | | | TTP: Ampsum (square root normalized) | | | CDA: Tonic Mean (square root normalized) | | |
|---|---|---|---|---|---|---|---|---|---|---|---|---|---|
| Normalization | *Predictors* | *Estimates* | *CI* | *p* | *Estimates* | *CI* | *p* | *Estimates* | *CI* | *p* | *Estimates* | *CI* | *p* |
| Square root normalized | (Intercept) | 0.75 | 0.59 – 0.90 | **<0.001** | 0.42 | 0.34 – 0.49 | **<0.001** | 0.55 | 0.42 – 0.68 | **<0.001** | 3.02 | 2.54 – 3.49 | **<0.001** |
| | Group | -0.53 | -0.90 – -0.16 | **0.005** | -0.26 | -0.44 – -0.08 | **0.004** | -0.39 | -0.69 – -0.09 | **0.01** | -0.15 | -1.29 – 0.99 | 0.797 |
| | Temperature | 0.44 | 0.30 – 0.58 | **<0.001** | 0.25 | 0.17 – 0.33 | **<0.001** | 0.49 | 0.33 – 0.64 | **<0.001** | 0 | -0.05 – 0.05 | 0.971 |
| | Trial | -0.03 | -0.04 – -0.03 | **<0.001** | -0.02 | -0.02 – -0.01 | **<0.001** | -0.03 | -0.04 – -0.02 | **<0.001** | -0.03 | -0.04 – -0.02 | **<0.001** |
| | Group * Temperature | -0.41 | -0.75 – -0.08 | **0.016** | -0.24 | -0.43 – -0.05 | **0.014** | -0.44 | -0.81 – -0.07 | **0.021** | -0.02 | -0.13 – 0.10 | 0.747 |
| | Group * Trial | 0.02 | -0.00 – 0.04 | 0.072 | 0.01 | 0.00 – 0.02 | **0.042** | 0.02 | 0.00 – 0.04 | **0.044** | 0 | -0.02 – 0.02 | 0.993 |
| | Temperature * Trial | -0.02 | -0.05 – -0.00 | **0.036** | -0.01 | -0.03 – -0.00 | **0.018** | -0.01 | -0.03 – 0.01 | 0.285 | -0.01 | -0.03 – 0.01 | 0.465 |
| | (Group * Temperature) * Trial | 0.02 | -0.04 – 0.07 | 0.577 | 0.01 | -0.02 – 0.04 | 0.6 | 0.01 | -0.04 – 0.06 | 0.691 | -0.01 | -0.06 – 0.04 | 0.763 |
| Square root normalized & Z-scored | (Intercept) | 0.01 | -0.04 – 0.06 | 0.688 | 0.01 | -0.04 – 0.06 | 0.688 | 0.01 | -0.04 – 0.05 | 0.826 | 0.01 | -0.04 – 0.06 | 0.613 |
| | Group | -0.01 | -0.12 – 0.11 | 0.918 | -0.01 | -0.12 – 0.11 | 0.924 | 0 | -0.12 – 0.11 | 0.939 | -0.01 | -0.13 – 0.10 | 0.807 |
| | Temperature | 0.85 | 0.61 – 1.10 | **<0.001** | 0.87 | 0.63 – 1.11 | **<0.001** | 0.93 | 0.68 – 1.18 | **<0.001** | 0.04 | -0.09 – 0.16 | 0.582 |
| | Trial | -0.08 | -0.10 – -0.06 | **<0.001** | -0.08 | -0.10 – -0.06 | **<0.001** | -0.06 | -0.08 – -0.04 | **<0.001** | -0.12 | -0.14 – -0.10 | **<0.001** |
| | Group * Temperature | -0.44 | -1.02 – 0.13 | 0.133 | -0.47 | -1.05 – 0.10 | 0.109 | -0.31 | -0.91 – 0.29 | 0.308 | 0 | -0.30 – 0.30 | 0.992 |

| | | | | | | | | | | | | | |
|---|---|---|---|---|---|---|---|---|---|---|---|---|---|
| | Group * Trial | -0.02 | -0.07 – 0.03 | 0.438 | -0.01 | -0.06 – 0.04 | 0.661 | -0.01 | -0.06 – 0.04 | 0.76 | 0.02 | -0.03 – 0.08 | 0.361 |
| | Temperature * Trial | -0.1 | -0.15 – -0.05 | **<0.001** | -0.1 | -0.15 – -0.05 | **<0.001** | -0.05 | -0.10 – 0.00 | 0.06 | -0.02 | -0.08 – 0.03 | 0.416 |
| | (Group * Temperature) * Trial | -0.01 | -0.13 – 0.11 | 0.839 | -0.02 | -0.14 – 0.10 | 0.717 | 0.01 | -0.11 – 0.13 | 0.849 | 0 | -0.13 – 0.12 | 0.963 |

[c]. This table presents results of linear mixed models predicting heat-evoked skin conductance response (SCR) as a function of Temperature (High vs Medium vs Low), Group (VMPFC vs Healthy Control), and Trial during the main experiment. Separate models were conducted for each SCR outcome measure derived from Ledalab's Continuous Decomposition Analysis (CDA) and trough-to-peak (TTP) analyses[7],

Supplementary Table S4.Cue effects on SCR.[d]

| Outcome measure | | CDA: Ampsum | | | CDA: SCR | | | TTP: Ampsum | | | CDA: Tonic Mean | | |
|---|---|---|---|---|---|---|---|---|---|---|---|---|---|
| Normalization | *Predictors* | *Estimates* | *CI* | *p* | *Estimates* | *CI* | *p* | *Estimates* | *CI* | *p* | *Estimates* | *CI* | *p* |
| Square root normalized | (Intercept) | 0.64 | 0.51 – 0.78 | **<0.001** | 0.36 | 0.30 – 0.43 | **<0.001** | 0.44 | 0.33 – 0.56 | **<0.001** | 3.01 | 2.54 – 3.49 | **<0.001** |
| | Group | -0.48 | -0.80 – -0.16 | **0.003** | -0.23 | -0.38 – -0.07 | **0.004** | -0.32 | -0.59 – -0.05 | **0.02** | -0.29 | -1.42 – 0.84 | 0.614 |
| | Cue | 0.07 | 0.00 – 0.13 | **0.047** | 0.04 | 0.01 – 0.07 | **0.024** | 0.08 | 0.02 – 0.14 | **0.013** | 0.01 | -0.04 – 0.05 | 0.797 |
| | Trial | -0.03 | -0.04 – -0.01 | **0.001** | -0.01 | -0.02 – -0.01 | **<0.001** | -0.02 | -0.04 – -0.01 | **0.003** | -0.02 | -0.03 – -0.01 | **<0.001** |
| | Group * Cue | -0.05 | -0.20 – 0.11 | 0.549 | -0.03 | -0.11 – 0.05 | 0.509 | -0.07 | -0.22 – 0.07 | 0.329 | -0.01 | -0.12 – 0.11 | 0.916 |
| | Group * Trial | 0.01 | -0.02 – 0.05 | 0.515 | 0.01 | -0.01 – 0.03 | 0.252 | 0.01 | -0.02 – 0.05 | 0.438 | 0 | -0.03 – 0.03 | 0.938 |
| | Cue * Trial | 0 | -0.03 – 0.03 | 0.804 | 0 | -0.02 – 0.02 | 0.982 | 0.01 | -0.02 – 0.04 | 0.373 | 0 | -0.02 – 0.02 | 0.942 |
| | (Group * Cue) * Trial | -0.01 | -0.08 – 0.07 | 0.892 | 0 | -0.04 – 0.04 | 0.93 | -0.02 | -0.09 – 0.05 | 0.515 | 0 | -0.06 – 0.05 | 0.917 |
| Square root normalized & Z-scored | (Intercept) | -0.22 | -0.29 – -0.14 | **<0.001** | -0.22 | -0.30 – -0.15 | **<0.001** | -0.22 | -0.30 – -0.13 | **<0.001** | 0.03 | -0.08 – 0.14 | 0.585 |
| | Group | -0.02 | -0.20 – 0.15 | 0.818 | 0 | -0.18 – 0.18 | 0.984 | 0.04 | -0.16 – 0.23 | 0.723 | -0.26 | -0.52 – -0.00 | **0.046** |
| | Cue | 0.13 | 0.00 – 0.26 | **0.047** | 0.13 | 0.01 – 0.26 | **0.037** | 0.16 | 0.03 – 0.28 | **0.016** | 0.02 | -0.13 – 0.17 | 0.829 |
| | Trial | -0.05 | -0.08 – -0.02 | **0.001** | -0.06 | -0.09 – -0.03 | **<0.001** | -0.05 | -0.08 – -0.02 | **0.002** | -0.12 | -0.15 – -0.08 | **<0.001** |

| | | | | | | | | | | | | | |
|---|---|---|---|---|---|---|---|---|---|---|---|---|---|
| | Group * Cue | 0.03 | -0.28 – 0.34 | 0.863 | 0.02 | -0.28 – 0.32 | 0.9 | 0.04 | -0.26 – 0.34 | 0.804 | 0 | -0.36 – 0.35 | 0.985 |
| | Group * Trial | -0.03 | -0.10 – 0.04 | 0.408 | 0 | -0.07 – 0.07 | 0.918 | -0.02 | -0.09 – 0.05 | 0.606 | 0.04 | -0.04 – 0.13 | 0.333 |
| | Cue * Trial | -0.02 | -0.09 – 0.04 | 0.449 | -0.02 | -0.08 – 0.04 | 0.608 | -0.01 | -0.07 – 0.05 | 0.832 | 0.03 | -0.04 – 0.10 | 0.409 |
| | (Group * Cue) * Trial | -0.06 | -0.21 – 0.09 | 0.431 | -0.04 | -0.18 – 0.11 | 0.61 | -0.09 | -0.23 – 0.05 | 0.228 | -0.01 | -0.18 – 0.16 | 0.869 |

[d]. This table presents results of linear mixed models predicting heat-evoked skin conductance response (SCR) on medium heat trials as a function of Cue (High vs Low), Group (VMPFC vs Healthy Control), and Trial during the main experiment. Separate models were conducted for each SCR outcome measure derived from Ledalab's Continuous Decomposition Analysis (CDA) and trough-to-peak (TTP) analyses[7],

Supplementary Table S5. Correspondence between SCR and pain rating on medium heat trials.[e]

| | | CDA: Ampsum | | | | | CDA: SCR | | | | | TTP: Ampsum | | | | | CDA: Tonic Mean | | | | |
|---|---|---|---|---|---|---|---|---|---|---|---|---|---|---|---|---|---|---|---|---|---|
| Rating type | | Estimate | Std. Error | df | t value | Pr(>\|t\|) | Estimate | Std. Error | df | t value | Pr(>\|t\|) | Estimate | Std. Error | df | t value | Pr(>\|t\|) | Estimate | Std. Error | df | t value | Pr(>\|t\|) |
| Intensity | (Intercept) | -0.192 | 0.035 | 20.02 | -5.508 | **0.0000** | -0.201 | 0.036 | 19.83 | -5.634 | **0.0000** | -0.191 | 0.041 | 19.53 | -4.656 | **0.0002** | 0.060 | 0.055 | 19.31 | 1.087 | 0.2905 |
| | Group | -0.022 | 0.082 | 19.58 | -0.27 | 0.7900 | -0.007 | 0.084 | 19.41 | -0.084 | 0.9336 | 0.036 | 0.097 | 19.19 | 0.373 | 0.7131 | -0.286 | 0.130 | 19.02 | -2.198 | **0.0406** |
| | Rating | 0.061 | 0.012 | 17.41 | 4.925 | **0.0001** | 0.057 | 0.011 | 15.88 | 5.051 | **0.0001** | 0.061 | 0.011 | 17.92 | 5.481 | **0.0000** | 0.063 | 0.015 | 16.91 | 4.058 | **0.0008** |
| | Group x Rating | -0.030 | 0.028 | 13.80 | -1.074 | 0.3012 | -0.030 | 0.025 | 12.06 | -1.193 | 0.2556 | -0.019 | 0.025 | 13.44 | -0.762 | 0.4593 | -0.044 | 0.035 | 13.76 | -1.258 | 0.2294 |
| Unpleasantness | (Intercept) | -0.168 | 0.037 | 19.61 | -4.538 | **0.0002** | -0.180 | 0.037 | 18.66 | -4.816 | **0.0001** | -0.170 | 0.041 | 18.79 | -4.132 | **0.0006** | 0.094 | 0.055 | 18.57 | 1.703 | 0.1052 |
| | Group | -0.043 | 0.087 | 18.51 | -0.499 | 0.6238 | -0.028 | 0.088 | 17.70 | -0.317 | 0.7552 | 0.012 | 0.097 | 17.95 | 0.122 | 0.9046 | -0.325 | 0.129 | 17.85 | -2.513 | **0.0218** |
| | Rating | 0.076 | 0.018 | 15.38 | 4.306 | **0.0006** | 0.069 | 0.016 | 14.57 | 4.285 | **0.0007** | 0.074 | 0.016 | 17.40 | 4.531 | **0.0003** | 0.090 | 0.025 | 19.16 | 3.579 | **0.0020** |
| | Group x Rating | -0.040 | 0.040 | 15.11 | -0.992 | 0.3370 | -0.039 | 0.037 | 14.48 | -1.064 | 0.3050 | -0.035 | 0.037 | 17.22 | -0.956 | 0.3525 | -0.071 | 0.057 | 18.33 | -1.228 | 0.2348 |

[e]. This table presents results of linear mixed models predicting heat-evoked skin conductance response (SCR) on medium heat trials as a function of Pain rating (mean-centered) and Group (VMPFC vs Healthy Control) during the experiment. Separate models were conducted for each SCR outcome measure derived from Ledalab's Continuous Decomposition Analysis (CDA) and trough-to-peak (TTP) analyses[7] and for each rating type.
